# Spatial Richness of Neural Magnetic Fields

**DOI:** 10.1101/2025.05.08.652049

**Authors:** Ziad Ali, Ada S. Y. Poon

**Author notes:** Senior Author (Ada S. Y. Poon).

## Abstract

Brain implants that measure neural magnetic fields, rather than electrical potentials, are expected to confer significant clinical advantages related to implant longevity and signal fidelity due to the elimination of the electrode-tissue interface. However, the informational differences between neural electrical potentials and magnetic fields remain poorly understood. Using a mathematical formalism based on neuronal current sources, we directly establish the complementary informational content of extracellular magnetic fields and electrical potentials. We then use computational modeling to illustrate how dense networks of neurons are easier to distinguish and spike sort on the basis of their magnetic, rather than electrical, spike templates. Lastly, we show how the solenoidal nature of neural magnetic fields facilitates approximate morphological reconstruction, even with sparse sensor arrays. Our findings highlight the unique experimental advantages of neural magnetic field sensing, motivating the development of compact, low-noise devices capable of meeting the stringent sensitivity requirements for single-shot cortical recordings.

## INTRODUCTION

Over the past two decades, researchers have achieved significant breakthroughs in decoding neural representations of cognitive intentions, such as limb movements^1^ and speech^2^, as well as external stimuli, including visual^3^ and auditory^4^ signals. These advances, which are foundational to the development of next-generation neuroprosthetics, critically depend on specialized hardware capable of precisely recording the activity of hundreds to thousands of densely packed neurons simultaneously. Implantable microelectrode arrays (MEAs) remain virtually the only clinically deployed technology capable of enabling such recordings, with devices such as the Utah array^5^ and Neuropixels probe^6^ widely adopted for single-cell resolution recordings in human and primate cortex.

Nonetheless, implantable MEAs face several significant challenges and limitations. A critical issue is the requirement for direct tissue contact, essential for providing conductive paths for current flow between the electrodes and target neurons. This requirement reduces the long-term efficacy of chronically implanted arrays because of gliosis, an immune response resulting in scar tissue formation around foreign bodies. Such scar tissue encapsulates the implant, impeding current flow^7^. For example, one study observed that the electrical potential amplitudes measured by implanted MEAs decreased by an average of 2.4% per month^8^. Consequently, implants often fail in less than a few years^1,9,10^. Organic microelectrodes, despite their inherently lower impedance, do not necessarily offer better performance, because tissue responses can substantially alter impedance irrespective of initial baseline values^11^. Moreover, the metal-solution interface of microelectrodes forms a capacitive double-layer effect, leading to charge screening that negatively affects signal quality^12^. Finally, MEAs can present practical challenges in *in vitro* settings because certain cell types exhibit poor adhesion to exposed metallic surfaces^13,14^.

From a device perspective, another major limitation of electrodes is the tradeoff between sensitivity and selectivity. Smaller electrodes improve spatial resolution, but their heightened electrical impedance results in worse thermal noise and signal loss^11^. In addition, since microelectrodes measure electrical potential differences, all arrays must include a reference electrode. Optimizing the size and placement of this electrode presents a non-trivial tradeoff: while a larger reference electrode offers lower impedance and thus reduced thermal noise, it does not measure common-mode signals identically, leading to imperfect noise filtering. Conversely, using a reference electrode identical in size to signal electrodes ensures better common-mode rejection, but introduces non-negligible thermal noise of its own^15^. Furthermore, placing the reference electrode too close to signal electrodes can unintentionally attenuate correlated neural signals^16^, while positioning it farther away increases the device footprint and risks introducing non-common-mode noise from other brain regions^17^.

Unlike electric field sensing, magnetic field sensing does not depend on a conductive path between sensor and tissue, making it a promising approach to overcome the aforementioned limitations. When neurons spike, they generate small magnetic fields in much the same way that current-carrying wires generate rotating magnetic fields according to Maxwell’s equations^18^. Since the magnetic permeability of tissue is essentially the same as air, these fields propagate undistorted throughout the brain and even outside the body, unaffected by the dielectric properties of tissue which smear and filter electric currents^19^. Magnetic sensors also measure absolute field values, eliminating the need for reference sensors, and can be fully encapsulated, thereby mitigating the sensitivity-selectivity tradeoff which primarily emerges from the impedance of the electrode-tissue interface^20^.

The deployment of magnetic sensing in neuroscience remains constrained by the extremely weak magnitude of neural magnetic fields—typically in the low picotesla to femtotesla range— relative to the noise floor of current microscale sensor technologies^21^. Existing demonstrations have primarily focused on obtaining low-SNR signals from large neurons or aggregated cellular activity, requiring extensive averaging to extract meaningful data^20,22,23^. Single-shot magnetic recordings at the level of individual cortical neurons have yet to be achieved. Despite recognized engineering benefits—including encapsulation, reference-free operation, and insensitivity to tissue impedance—the pace of development has been slow, in part due to limited understanding of the distinct *information* that magnetic sensing can provide over traditional electrical recordings.

In this work, we present a rigorous mathematical formulation describing how spiking neurons generate extracellular electrical potentials and magnetic fields, showing that these signals arise from fundamentally distinct current sources: transmembrane currents for electric fields and longitudinal currents for magnetic fields. From this distinction, three key insights follow: electric and magnetic fields convey complementary information about the neuron and its neural activity; they scale differently with distance from the source; and they exhibit distinct vector field properties— curl-free for electric fields versus divergence-free for magnetic fields. Building on these insights, we investigate how magnetic and electric signals differ in their ability to distinguish closely spaced neurons using only extracellular spike waveforms. We quantify this difference in distinguishability by comparing pairwise spatial resolution and network-wide channel conditioning across large neural populations. To assess practical implications, we apply spike sorting to simulated populations of both *in vitro* (planar) and *in vivo* (three-dimensional) neurons. Lastly, we demonstrate how the solenoidal nature of magnetic fields enables new approaches to neural morphology reconstruction using off-the-shelf algorithms.

## RESULTS

### Complementary Information from Extracellular Neural Fields

A neuron can be conceptually divided into three regions: the intracellular space, the membrane region, and the extracellular space Fig. 1a). By assuming that all non-linear sources are confined to the membrane region, the neural fields in the intracellular and extracellular spaces satisfy the Poisson equation. Consequently, Green’s theorem can be applied to these spaces^24^. Previous studies have applied Green’s theorem to express the extracellular potential in terms of the current densities and potentials on the inner and outer surfaces of the cell membrane, or, alternatively, in terms of the transmembrane potential^25–27^. Here, we extend this approach using the vector form of Green’s theorem to relate the extracellular magnetic field to the current density on the inner surface of the cell membrane. This derivation aims to succinctly show how the extracellular magnetic field and electrical potential arise from distinct components of this current density.

**Figure 1.**
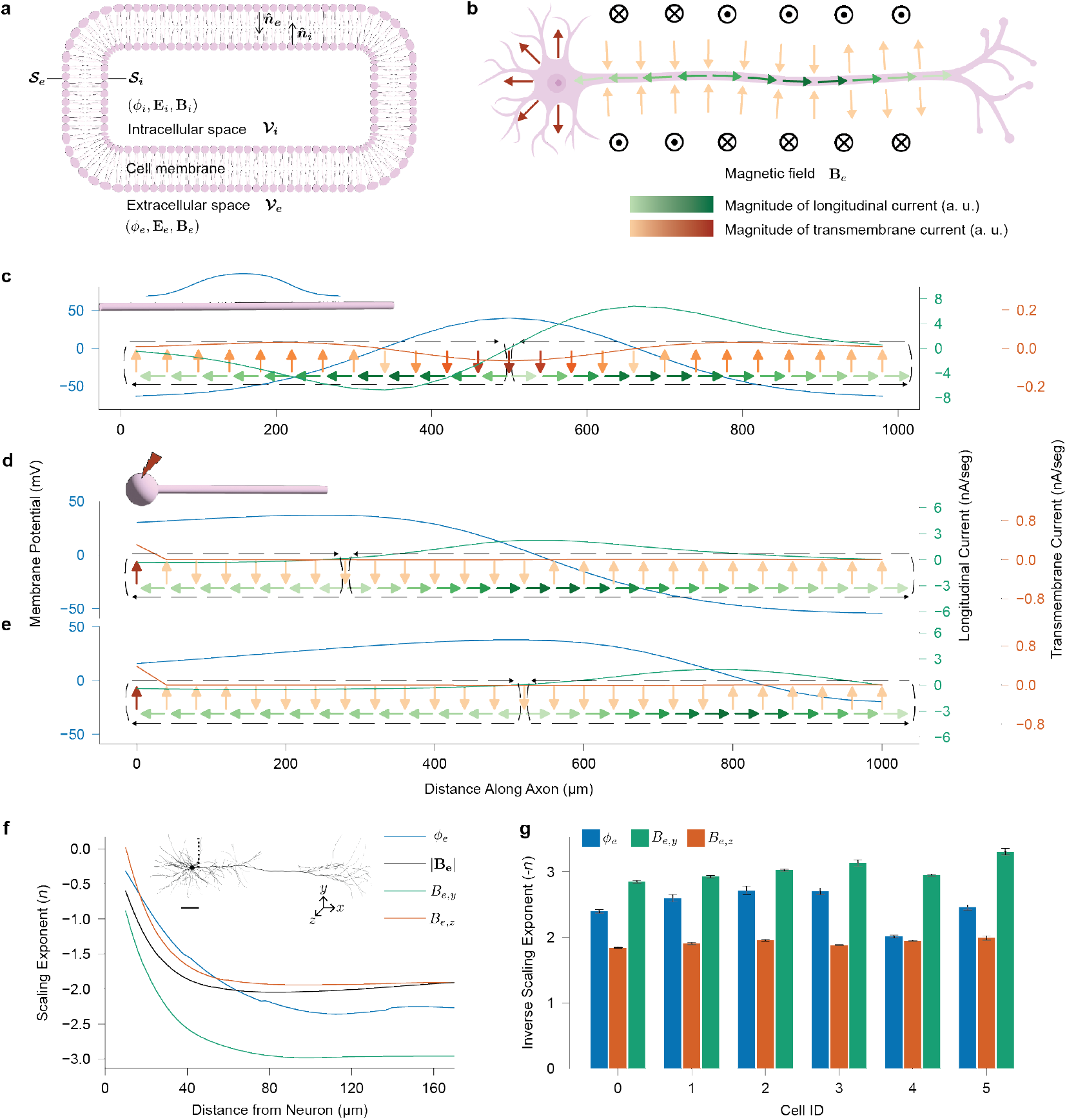
Application of Green’s theorem and scaling analysis of intracellular currents and extracellular fields in neuron models. **a**, Application of Green’s theorem to the intra-cellular space, 𝒱_*i*_, of a neuron—characterized by the potential *ϕ*_*i*_, electric field **E**_**i**_, and magnetic field **B**_**i**_, bounded by the surface 𝒮_*i*_ with local normal 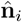—and to the extracellular space, 𝒱_*e*_—characterized by the potential *ϕ*_*e*_, electric field **E**_**e**_, and magnetic field **B**_**e**_, bounded by the surface 𝒮_*e*_ with local normal 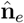. **b**, Illustration of longitudinal and transmembrane currents generated by a neuron during an action potential, along with the direction of the extracellular magnetic field induced by longitudinal currents. **c**, Analytically-derived distribution of longitudinal 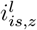 and transmembrane 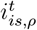 components of the intracellular surface current along an infinitely extended cylindrical axon with a Gaussian-distributed membrane potential *ϕ*_*m*_, characterized by a standard deviation of 167 *µ*m. The black dashed lines indicate current loops composed of the transmembrane and longitudinal currents which propagate down the length of the axon during the action potential. **d, e**, Cellular current distributions for a ball-stick cell model when the action potential peak travels (**d**) one quarter and (**e**) half of the axon length. Parameters: axon length = 1 mm; axon diameter = 2 *µ*m; soma diameter = 20 *µ*m; segment length = 0.5 *µ*m. **f**, Scaling exponent *n* characterizing how the extracellular potential *ϕ*_*e*_ and magnetic field **B**_**e**_, including the planar (*B*_*e*,*y*_) and normal (*B*_*e*,*z*_) components relative to the sensor array plane, vary as a function of distance from the cell (*R*^*n*^). The upper part of the figure shows the cell model (Cell 2 from SFig. 1) and measurement locations used to evaluate the scaling, marked by the black dashed line. Scale bar, 100 *µ*m. **g**, Asymptotic negative scaling exponent *− n* for the extracellular potential *ϕ*_*e*_ and the magnetic field components *B*_*e*,*y*_ and *B*_*e*,*z*_ for cells 0–5 (SFig. 1). Evaluated between 0 to 50 *µ*m down the length of the apical dendrite, *N* = 52.

The vector form of Green’s theorem states that

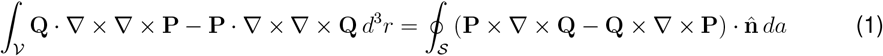

for any well-behaved vector fields **P** and **Q** defined in the volume 𝒱 enclosed by the surface 𝒮^28^. Now, we choose **P** = **H**, the magnetic field intensity, and **Q** = **c***/* |**r***−* **r**^*′*^| for any constant vector **c**, where **r** is the observation point and **r**^*′*^ is the integration variable. Equation (1) becomes

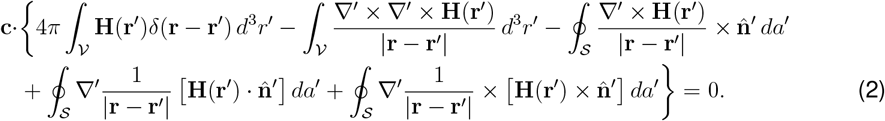

The detailed proof is included in the supplementary information. Since (2) is true for all **c**, the expression inside the curly brackets should be zero, yielding our desired identity.

Referring to Fig. 1a, we apply this identify to the combined volume 𝒱= 𝒱_*e*_ *∪* 𝒱_*i*_. Given that the membrane is very thin, 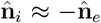 denoted simply by 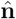 and 𝒮_*i*_ ≈ 𝒮_*e*_ denoted by 𝒮. Additionally, we utilize the relation *∇ ×* **H** = **J**_*imp*_ *− σ ∇ ϕ* where **J**_*imp*_ is the impressed source, *ϕ* is the electrical potential, and *σ* is the conductivity. The impressed source is confined entirely within the volume, with no contribution on the surface. Consequently, for all **r** *∈* 𝒱_*e*_,

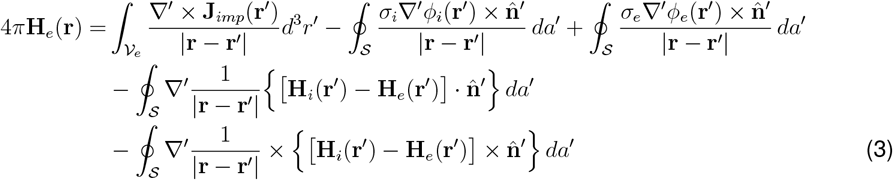

where the subscripts *e* and *i* denote variables in the extracellular and intracellular spaces, respectively. Due to the membrane’s much lower conductivity compared to both *σ*_*e*_ and *σ*_*i*_, the longitudinal current within the membrane is negligible. Combined with the membrane’s small thickness, this results in the magnetic field remaining nearly continuous across the membrane, rendering the last two integrals negligible. Thus, the extracellular magnetic field can be described as the sum of the field originated by the solenoidal component of the impressed source (for example, magnetic stimulation from a microcoil) in an unbounded medium and the field originated by the longitudinal component (orthogonal to 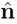) of the current density along the inner and outer surfaces of the cell membrane.

Similarly, by applying the scalar form of Green’s second identity to 𝒱 = 𝒱_*e*_, for all **r** *∈* 𝒱_*e*_ ^27^,

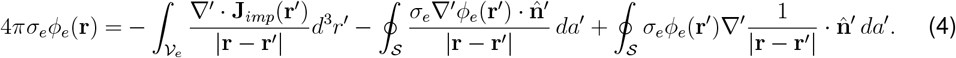

Given the membrane’s small thickness and its conductivity being much lower than both *σ*_*e*_ and *σ*_*i*_, the transverse current is approximately continuous across the membrane, leading to 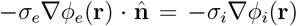 on *𝒮*. As a result, the extracellular potential can be described as the sum of the field originated by the irrotational component of the impressed source (for example, electrical stimulation from microelectrodes) in an unbounded medium and the field originated by the transverse component (parallel to 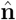) of the current density crossing the inner or outer surfaces of the cell membrane.

To highlight how the extracellular magnetic field and electrical potential arise from distinct components of the current density on the inner surface of the membrane defined by **i**_*is*_ = *−σ*_*i*_ *∇ ϕ*_*i*_ on 𝒮_*i*_, we simplify by assuming a source-free and extended extracellular space. In such a space, the longitudinal component of the current density along the outer membrane surface and the extracellular potential at the outer membrane surface become negligible. Furthermore, assuming this space has negligible magnetic properties (magnetization **M** *≈* 0), we can linearly relate the magnetic flux density **B** to the magnetic field intensity by **B** = *µ*_0_**H**. Consequently,

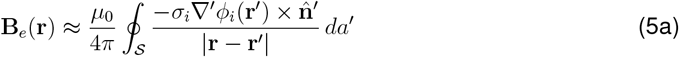

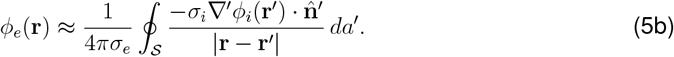

These expressions highlight a fundamental duality between the extracellular magnetic field and the electrical potential generated by the cell. The magnetic field arises from the longitudinal component of the intracellular surface current density, 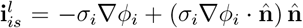 on *𝒮*_*i*_, while the electrical potential is determined by the transmembrane component, 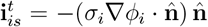 on 𝒮_*i*_. This complementary relationship means that magnetic field measurements primarily reveal currents flowing parallel to the membrane, while electrical potentials capture those crossing it. Together, they reveal orthogonal aspects of the neuron’s physical structure and underlying neural dynamics. We therefore hypothesize that simultaneous recordings of neural electrical potentials and magnetic fields can significantly enhance our ability to analyze the behavior and properties of large populations of neurons.

We further propose that, for certain cell types, neural magnetic fields may provide richer information than electrical potentials. This is because transmembrane currents, primarily generated at the soma, are the main contributors to electrical potentials, while longitudinal currents along extended processes, such as axons and apical dendrites, are the principal sources of magnetic fields. Given the greater morphological complexity of these processes, we predict that neurons with extended, linear regions—such as cortical neurons—can be more readily distinguished by their extracellular magnetic fields than by their electrical potentials. This distinction is further highlighted by the rotational pattern of the magnetic field around long processes, producing opposing polarities at different spatial locations relative to the cell (see Fig. 1b). In contrast, transmembrane currents from the soma yield extracellular electrical potentials with nearly uniform polarity, resulting in less spatial variation. We therefore hypothesize that the sensitivity of neural magnetic fields to long processes will enhance our ability to estimate and reconstruct cellular morphologies. To investigate how these theoretical distinctions manifest in realistic measurements, we will analyze and model the extracellular magnetic fields and electrical potentials of various cell types in the following sections.

### Neural Signal Scaling

The scaling of extracellular potential with distance has been extensively studied^29–32^, but a similar understanding for the extracellular magnetic field remains relatively unexplored. Having established that the extracellular magnetic field and electrical potential can be expressed in terms of the orthogonal components 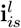 and 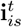, respectively, which together form the current density on the inner membrane surface, 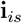, we now focus on how these components flip signs along the neuron. The frequency of these sign reversals determines the order of the resulting multipole fields, providing key insights into how signals scale with distance.

To focus on the sign reversal, we model the neuron as an infinitely extended cylindrical axon within a vast volume conductor. Solutions to the Laplace’s equation in cylindrical coordinates are given by the modified Bessel functions^33^. The intracellular and extracellular potentials can then be expressed as

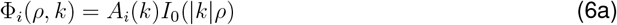

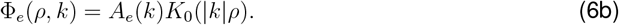

Here, Φ_*i/e*_(*ρ, k*) denotes the Fourier Transform of *ϕ*_*i/e*_(*ρ, z*) defined as 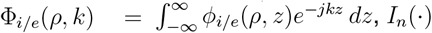 and *K*_*n*_(*·*) are modified Bessel functions of the first and second kind of order *n*, respectively, and *A*_*i/e*_(*k*)’s are functions determined by boundary conditions. To capture the neuron’s activity during firing, we follow the approach outlined in Refs. 34–36. We determine *A*_*i/e*_(*k*)’s such that solutions to the Laplace’s equation satisfy a given transmembrane potential, *ϕ*_*m*_(*z*), representing the action potential for an active neuron. As a result, the intracellular potential in the Fourier domain is given by

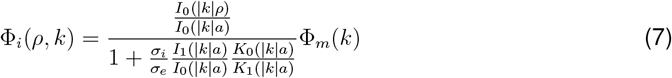

where *a* is the radius of the cylindrical axon. In the past, when computing power was limited, formulas derived from this approach provided a method for quantitatively evaluating fields from an active nerve. Nowadays, with vastly improved computing power and the emergence of advanced software libraries such as NEURON^37^ and LFPy^38^, these limitations no longer apply. However, here we utilize (7) to perform an analytical analysis of extracellular field scaling.

Given that the length of the axon is much greater than its radius, the characteristic length scale over which the transmembrane potential has significant values, denoted by *L*, is expected to be much larger than the radius of the axon, that is, *L≫ a*. As a result, the Fourier transform of the transmembrane potential, Φ_*m*_(*k*), becomes negligible for |*k*| greater than a few mulitples of 1*/L*. Thus, it is reasonable to assume that |*k*| *a≪* 1 and the modified Bessel functions can be approximated as follows:

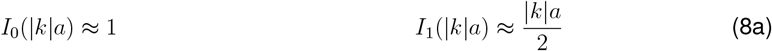

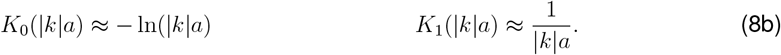

Applying these approximations, Equation (7) simplifies to

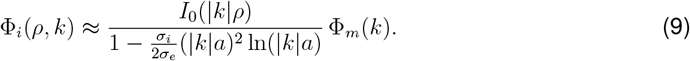

Since lim_*x→*0_ *x*^2^ ln *x* = 0, this reduces further to

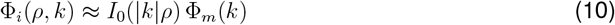

which yields the current components:

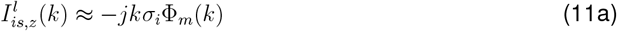

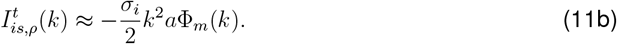

The proof is included in the supplementary information. Their inverse Fourier transforms are

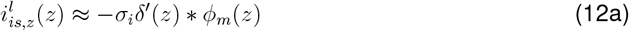

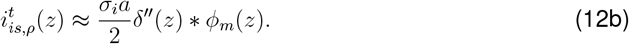

When *ϕ*_*m*_(*z*) is a unimodal function such as a Gaussian function, the longitudinal component, 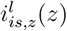, reverses sign once whereas the transmembrane component, 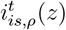 reverses sign twice, as illustrated in Fig.1b. Consequently, the extracellular magnetic field behaves like a dipole field while the electrical potential exhibits quadrupole characteristics. This implies that the extra-cellular magnetic field scales with distance *R* as 1*/R*^3^, while the extracellular electric field and potential scale as 1*/R*^4^ and 1*/R*^3^, respectively. Due to this latter quadrupole scaling, the extracellular electric potential measured by electrodes with a local reference is expected to scale between 1*/R*^4^ and 1*/R*^3^, that is 1*/R*^3+*ϵ*^ for some 0 *≤ ϵ ≤* 1.

Further derivation yields the extracellular fields:

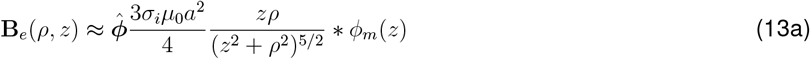

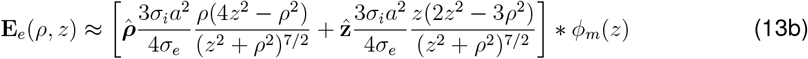

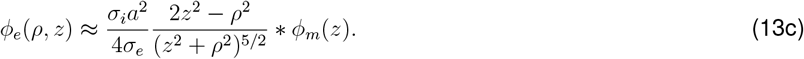

The proof is included in the supplementary information. For large distances, *z ~ ρ ~ R ≫* 1, the magnetic field scales as 1*/R*^3^ while the electric field and the potential scale as 1*/R*^4^ and 1*/R*^3^, respectively. In conclusion, for a cylindrical axon of infinite extent, the extracellular magnetic field exhibits scaling behavior comparable to, or better than, the electrical potential.

We now proceed with three numerical evaluations. First, using a Gaussian function for *ϕ*_*m*_(*z*), Fig. 1c shows the distribution of 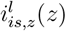 and 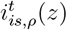 along a cylindrical axon, as derived from (12). These currents form two current loops, *n*, (black dashed arrows in Fig. 1c), corresponding to traveling waves of depolarization, which indicate the number of sign flips for the longitudinal (*n*) and transmembrane (*n* + 1) current components. These results confirm the dipole behavior of the magnetic field and the quadrupole behavior of the electrical potential.

Next, we implement a ball-stick model to compute the corresponding transmembrane and longitudinal currents. Figures 1d, e show the current distributions at the times when the action potential peaks at one-quarter and one-half of the length of the axon, respectively. Unlike a cylindrical axon, the presence of the soma significantly amplifies the transmembrane current at the proximal end relative to the rest of the axon. As a result, the quadrupole effect on the electric potential is reduced, suggesting that the scaling behavior for the extracellular electric potential may be better than 1*/R*^3+*ϵ*^. Similarly, the forward longitudinal current in the axon is stronger than the reverse, indicating that the dipole effect on the magnetic field is diminished, and its scaling behavior may be better than 1*/R*^3^.

To evaluate how the scaling behavior of actual neurons deviates from the ideal case, we simulate six morphologically realistic neurons (SFig. 1). These neurons are all layer V rat somatosensory cortical neurons provided by the Blue Brain Project^39^. They were chosen due to their highly detailed morphological and biophysical characteristics as well as their similarity to the information-rich neurons commonly targeted in human studies. The selected models include at least one of each pyramidal neuron type available in the Blue Brain Project’s layer V repository (TTPC1, TTPC2, UTPC, STPC) and two interneuron types (DBC, MC).

Figure 1f presents the scaling exponent *n* of the extracellular electrical potential (*ϕ*_*e*_) and magnetic field (**B**_*e*_) calculated along a radial line of measurement points extending from near the soma and adjacent to the apical dendrite of Cell 2 from SFig. 1 (Fig. 1f, inset). We also present the scaling exponents for the planar (*B*_*y*_) and normal (*B*_*z*_) components of **B**_*e*_, recognizing that most magnetic sensors measure magnetic fields along only a single axis. Our simulations reveal that the asymmetric distribution of transmembrane and longitudinal currents causes both the electrical potential and magnetic field to deviate from the expected 1*/R*^3^ scaling relationship. This deviation is more pronounced for the magnetic field, which scales approximately as 1*/R*^2^ at large distance, compared to the electrical potential, which scales approximately as 1*/R*^2.5^. Notably, the two vector components of the magnetic field exhibit distinct scaling behaviors: the planar component *B*_*y*_ decreases approximately an order of magnitude faster than the normal component *B*_*z*_. This discrepancy arises from the geometrical arrangement of the neuron relative to the measurement points. In conventional *in vitro* experiments, neurons lies flat above a planar array of sensors. If the sensor array is positioned a small distance Δ*z* below the neuron, for *R≫* Δ*z* the sensors are essentially coplanar with the neuron and can therefore only detect the normal component of the magnetic field around a major process, rendering the planar component negligible.

Figure 1g presents results of the scaling analysis across all analyzed cell morphologies. As shown, the normal magnetic field *B*_*z*_ consistently scales approximately with 1*/R*^2^, whereas the electrical potential *ϕ*_*e*_ scales close to 1*/R*^2.5^ in most cases. This consistent scaling behavior suggests that for *in vitro* measurements where sensor resolution is often limited, it may be more advantageous to measure the normal magnetic fields as they retain greater signal strength over a broader spatial range than electrical potentials.

### Cellular Spatial Resolution Limit

As elucidated in the complementary information section, when a biologically realistic neuron fires, it generates three-dimensional electric and magnetic vector fields uniquely shaped by its morphological distribution of transmembrane and longitudinal currents. Because the spatial distribution of these currents is unique to each neuron in a given population, each neuron has its own distinctive electric and magnetic field signatures. This uniqueness is crucial for identifying and distinguishing neurons solely on the basis of extracellular recordings. However, during an action potential, sensor arrays typically capture only two-dimensional (2D) snapshots of these fields, known as spike templates. This reduction in dimensionality complicates the differentiation of neurons that are spatially proximate and/or morphologically similar. In this section, we investigate how the neural signal type—electric or magnetic field—affects the minimum intercellular distance required for reliable pairwise cell discrimination based on 2D spike templates.

We analyze the six previously described layer V neuron models by generating simulated spike templates for both the extracellular electrical potential (*ϕ*_*e*_) and the three Cartesian components of the magnetic field (*B*_*e*,*x*_, *B*_*e*,*y*_, and *B*_*e*,*z*_). For brevity, we refer to the extracellular potential simply as *ϕ* and the magnetic field components as *B*_*x*_, *B*_*y*_, and *B*_*z*_, since our analysis is confined to the extracellular space. We define the base templates—referred to as spike signatures—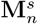, for neuron *n* and signal *s ∈ {ϕ, B*_*x*_, *B*_*y*_, *B*_*z*_*}*, as *N*_*R*_ *× N*_*T*_ matrices, where *N*_*R*_ denotes the number of spatial measurement points (sensors) and *N*_*T*_ is the number of time steps over the course of a single spike event. In our simulations, each neuron is positioned at the center of a dense grid of measurement points, with its major process oriented along the positive *x*-axis. The grid consists of 1,500 *×* 1,500 measurement points with a 2-*µ*m pitch, effectively approximating a continuous measurement space with near infinitesimal spatial resolution. This setup allows us to isolate the effect of neuronal morphology and field patterns, independent of constraints imposed by real-world sensor spacing.

We investigate how spike signatures from two neurons, separated by a spatial displacement Δ**r** = (Δ*x*, Δ*y*), may appear similar due to spatial proximity and orientation. To simulate this effect, we generate new templates from the base template by applying spatial transformations: a rotation by angle *θ* and a translation by Δ**r** (Fig. 2a). These transformed templates are denoted as 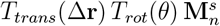, where *T*_*rot*_(*θ*) and *T*_*trans*_(**r**) represent the respective rotation and translation operators. To quantify the minimum intercellular distance, we introduce a cosine similarity metric that measures the similarity between the spike signature of neuron *n* and that of a spatially transformed neuron *m* for signal *s*:

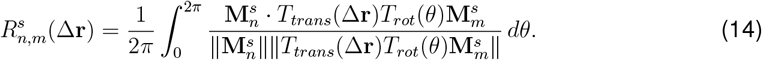

**Figure 2.**
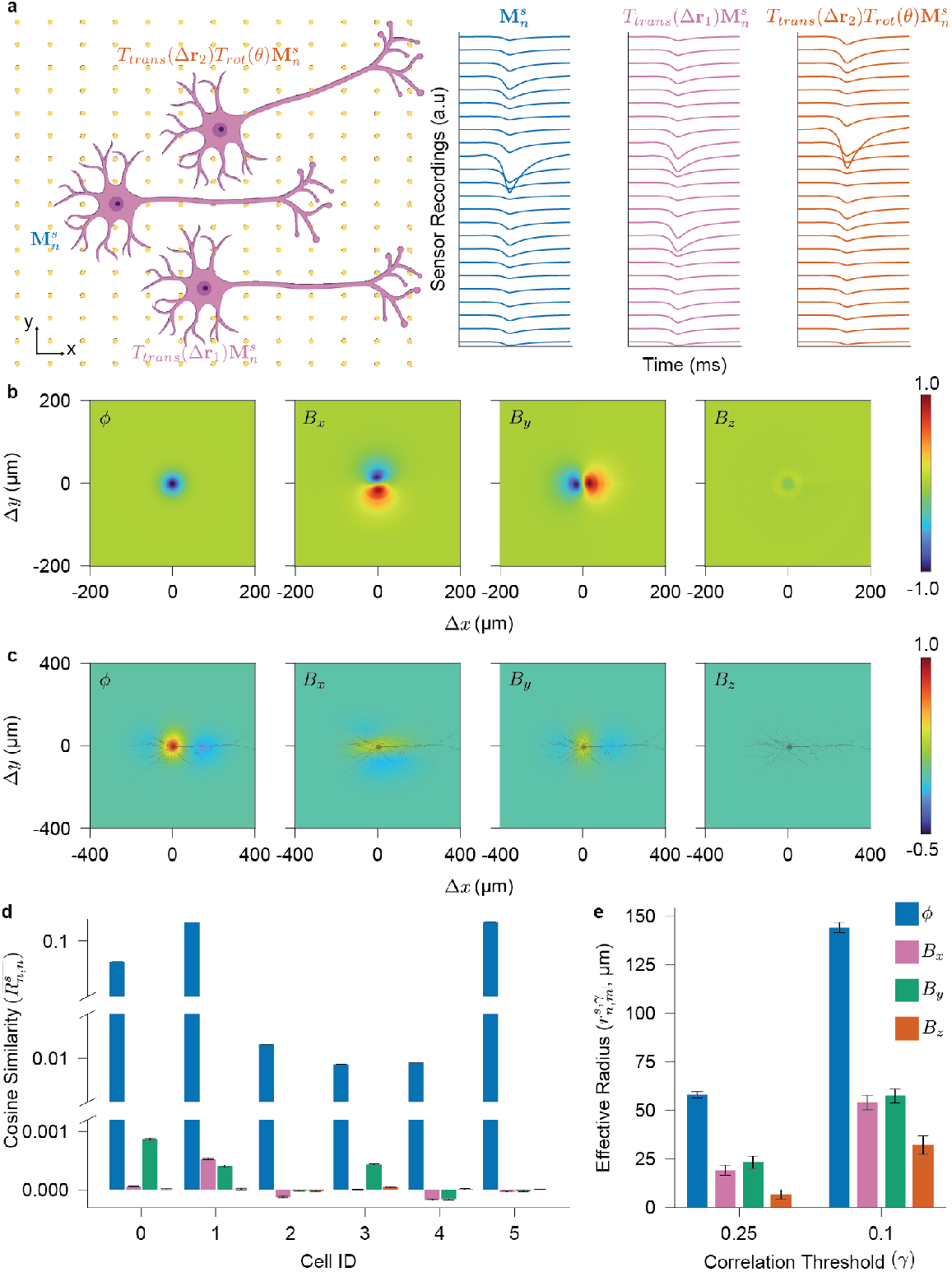
Spatial resolution limits of simulated neural extracellular electrical potentials and magnetic fields. **a**, Analysis setup and example spike templates. Neurons are positioned above a dense, simulated array of sensors. Spike templates are generated by translating, 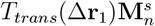, and rotating, 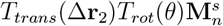, the neurons within the sensor plane. Displayed are abridged spike templates comprising 24 measurement points for three distinct neuron positions and orientations. **b**, Spread templates 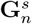 for cell *n* = 2 and signals *s ∈ {ϕ, B*_*x*_, *B*_*y*_, *B*_*z*_*}*. **c**, Spatial similarity 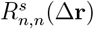 for cell *n* = 2 and signals *s ∈ {ϕ, B*_*x*_, *B*_*y*_, *B*_*z*_*}*. **d**, Mean spatial auto-similarity for Cells 0–5 from SFig. 1, averaged over displacements along both *x*- and *y*-axes within *±* 400 *µ*m. **e**, Effective radius for high (*γ >* 0.25) and moderate (*γ >* 0.1) similarity thresholds, averaged over all cell pairs (*n, m*) from SFig. 1.

By averaging over all rotation angles, we account for variations in neuronal orientation. A high value of 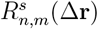, approaching 1, indicates that the spike signatures of neurons *n* and *m* are highly similar for signal type *s* at the displacement Δ**r**, suggesting that the two neurons produce nearly indistinguishable extracellular fields and are therefore difficult to resolve based on that signal type. Conversely, a low or negative 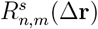 signifies a high degree of dissimilarity between the spike signatures, indicating that the neurons can be more easily distinguished at that displacement.

Under the assumption of infinitesimal pitch, the translation operator *T*_*trans*_(Δ**r**) can be factored outside the integral, and the rotation integral interpreted as an orientation-averaging operator *T*_*sweep*_, yielding

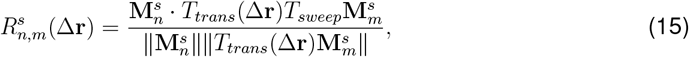

where

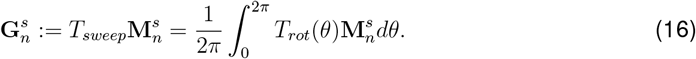

We refer to 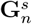 as the *spread template*. While its derivation differs from the classical point-spread function (PSF) in optics^40^, the spread template similarly captures the average spatial distribution of a neuron’s extracellular field at a given location. By serving as an analogue of a PSF, 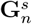 characterizes how neuronal signals disperse through space, providing a basis for evaluating neuronal distinguishability based solely on their spatial separation.

Figure 2b displays the spread templates for Cell 2 at the time corresponding to the maximum overall template signal power. For magnetic field templates, the rotation transformation is applied directly to the full vector field **B**_*e*_. Consequently, the vector-valued spread template is invariant to orientation. However, the individual components—specifically *B*_*x*_ and *B*_*y*_—are not, as they depend on how the rotated field projects onto their respective sensing axes, as illustrated in Fig. 2b. For visualization, the electrical potential template is normalized to its maximum value, while the magnetic field components are scaled by the maximum value across all three components to ensure consistent relative amplitudes. This normalization results in a lower relative magnitude for the *B*_*z*_ component compared to *B*_*x*_ and *B*_*y*_.

Figure 2c shows the auto-similarity, 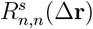 for cell *n* = 2, evaluated for each signal type *s ∈ {ϕ, B*_*x*_, *B*_*y*_, *B*_*z*_*}*. The electrical potential exhibits high similarity with a radially symmetric distribution centered at the soma—consistent with the complementary information section, where we identified the soma as the primary source of transmembrane currents generating uniform-polarity extracellular signals. This symmetry is also evident in the spread template for *ϕ* (Fig. 2b), indicating that neurons positioned around and near the soma produce highly similar extracellular potentials. In contrast, magnetic fields show lower spatial similarities. The planar magnetic fields (*B*_*x*_, *B*_*y*_) exhibit antisymmetric spread templates with respect to their sensing axes (Fig. 2b), due to their sensitivity to neuronal orientation. For instance, rotating a neuron by 180^*°*^ reverses the *B*_*y*_ signal, flipping the spread template across the *y*-axis. This leads to lower similarity and a triphasic spread template pattern slightly elongated along the sensing axis. The normal magnetic field (*B*_*z*_) is even more sensitive to rotation and translation. Sensors on opposite sides of the neuron’s extended structure detect signals of opposite polarity (Fig. 1b), and even small rotations around the soma or translations perpendicular to the neuron’s major process can render the *B*_*z*_ template nearly anti-similar with the original. Consequently, the normal magnetic field results in minimal spatial similarity, substantially lowering the separation limit for distinguishing neurons—outperforming planar magnetic fields and, even more so, electrical potentials.

Figure 2d presents the spatial similarity, averaged over displacements along both the *x* and *y* axes within a range of *±*400 *µ*m, for all six morphologically realistic neurons shown in SFig. 1. On average, electrical potential templates exhibit similarity values several orders of magnitude higher than those of magnetic field templates—even for cells *n* = 2, 3, 4, where extensive dendritic branching reduces radial symmetry. Planar magnetic field templates occasionally show non-negligible similarity or even anti-similarity, due to their antisymmetric spread templates and sensitivity to neuronal orientation. In contrast, normal magnetic field templates consistently exhibit near-zero similarity, making them particularly well-suited for recording and analyzing densely packed neuronal populations.

Finally, we quantify the spatial resolution limit—the minimum intercellular distance required for reliable discrimination—by defining an effective radius 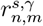 as the radius of a circle with an area equal to the region where the similarity 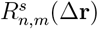 exceeds a threshold *γ*:

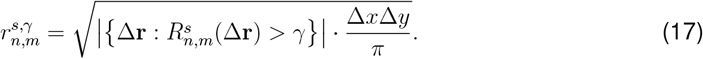

Here, |*·*| denotes the cardinality of the set. Figure 2d shows the effective radii averaged across all cell pairs (*n, m*) for two thresholds: 0.25 (high similarity) and 0.1 (moderate similarity). As expected, magnetic fields achieve higher spatial resolution than electrical potentials. Planar magnetic field templates yield effective radii approximately three times smaller than those of electrical potentials, while normal magnetic fields provide even greater resolution—up to five- to nine-fold improvements. These results indicate that magnetic sensors, especially those measuring the normal field, are fundamentally better suited for distinguishing neurons in densely populated regions than traditional electrical recordings.

### Multi-Source Discrimination in Practical Sensor Arrays

Having established that the spatial distributions of electrical potentials and magnetic fields fundamentally limit pairwise neuronal distinguishability, we now extend our analysis to large-scale neural networks, key for understanding higher-order cognitive functions. In practice, data throughput and hardware constraints limit the number of sensors that can be deployed in experimental or clinical arrays^41^. Consequently, achieving broader coverage to monitor more neurons necessitates sparser sensor placement—in contrast to the near-infinitesimal pitch in the previous section—which in turn reduces the spatial resolution and degrades the measurement quality for individual neurons. To evaluate how signal modality affects this trade-off, we adopt a metric from communication theory—the condition number—to quantify network-wise neuronal distinguishability. In addition to analyzing individual field components, we also examine combinations, such as (*ϕ, B*_*z*_), to assess whether integrating complementary electrical and magnetic recordings improves performance. In these multimodal configurations, each measurement location includes multiple sensors capable of recording different fields.

We adopt the condition number from communication theory, where it serves as a standard metric for signal separability in systems such as massive MIMO and spread-spectrum CDMA^42,43^. In MIMO systems, the channel matrix **H** encodes the coupling between transmit and receive antennas, and its condition number *κ*(**H**)—the ratio of the largest to smallest singular values—quantifies how easily individual data streams can be separated. In spread-spectrum CDMA (code division multiple access), multiple users transmit concurrently using distinct spreading codes, aiming to differentiate as many users as possible despite the codes not being strictly orthogonal. In both cases, the condition number reflects how well overlapping signals can be separated, with a value close to 1 indicating near-orthogonality, and a value much greater than 1 indicating substantial challenges in signal separation, especially under noisy conditions. Drawing on this analogy, we use the condition number to quantify how effectively different extracellular signal modalities support neuronal distinguishability in densely populated networks.

In our analysis, the channel matrix is constructed by flattening each neuron’s simulated spike template—originally an *N*_*R*_ *× N*_*T*_ matrix—into an *N*_*R*_*N*_*T*_ *×* 1 column vector. Vectors from *N*_*C*_ randomly sampled cells, each with a different rotation and translation, are concatenated to form an *N*_*C*_ *×N*_*R*_*N*_*T*_ channel matrix. This construction assumes simultaneous firing of all neurons under test, reflecting scenarios such as bursting activity or high synchrony. While idealized, this assumption serves as a conservative estimate of channel performance under the most challenging conditions for neuronal separability.

We use the condition number to evaluate neuronal distinguishability in two experimental scenarios. First, we fix the number of sensors and vary the sensor pitch—50, 75, and 100 *µ*m—in a 10 *×* 10 array to examine how increasing the coverage area affects distinguishability, while maintaining a constant cell density of 340 cells/mm^2 44^. Second, we fix the sensor pitch at 75 *µ*m— thereby holding the coverage area constant—and vary the number of neurons from 20 to 200 to assess the effect of increasing cell density on neuronal separability.

Figure 3a presents the trade-off between neuronal distinguishability—measured as the inverse of the condition number—and array coverage using channel matrices constructed from the six morphologically realistic neurons shown in SFig. 1. Combining electrical potential and normal magnetic field measurements (*ϕ, B*_*z*_) achieves the optimal trade-off, supporting our hypothesis that these signals provide complementary information. In contrast, arrays measuring only a single planar magnetic field component (*B*_*x*_ or *B*_*y*_) yield the poorest performance, contradicting the results observed in the pairwise distinguishability analysis. This discrepancy arises because neuronal processes aligned with the sensing axis generate magnetic fields orthogonal to it (via the right-hand rule), making such neurons nearly undetectable. Measuring both planar components (*B*_*x*_, *B*_*y*_) mitigates this orientation sensitivity and yields performance comparable to the *B*_*z*_-only array, which is inherently less susceptible to orientation-dependent signal loss on a planar array. Simultaneous measurement of all three magnetic components (*B*_*x*_, *B*_*y*_, *B*_*z*_) captures additional information and further improves performance, closely approaching that of the hybrid (*ϕ, B*_*z*_) array.

**Figure 3.**
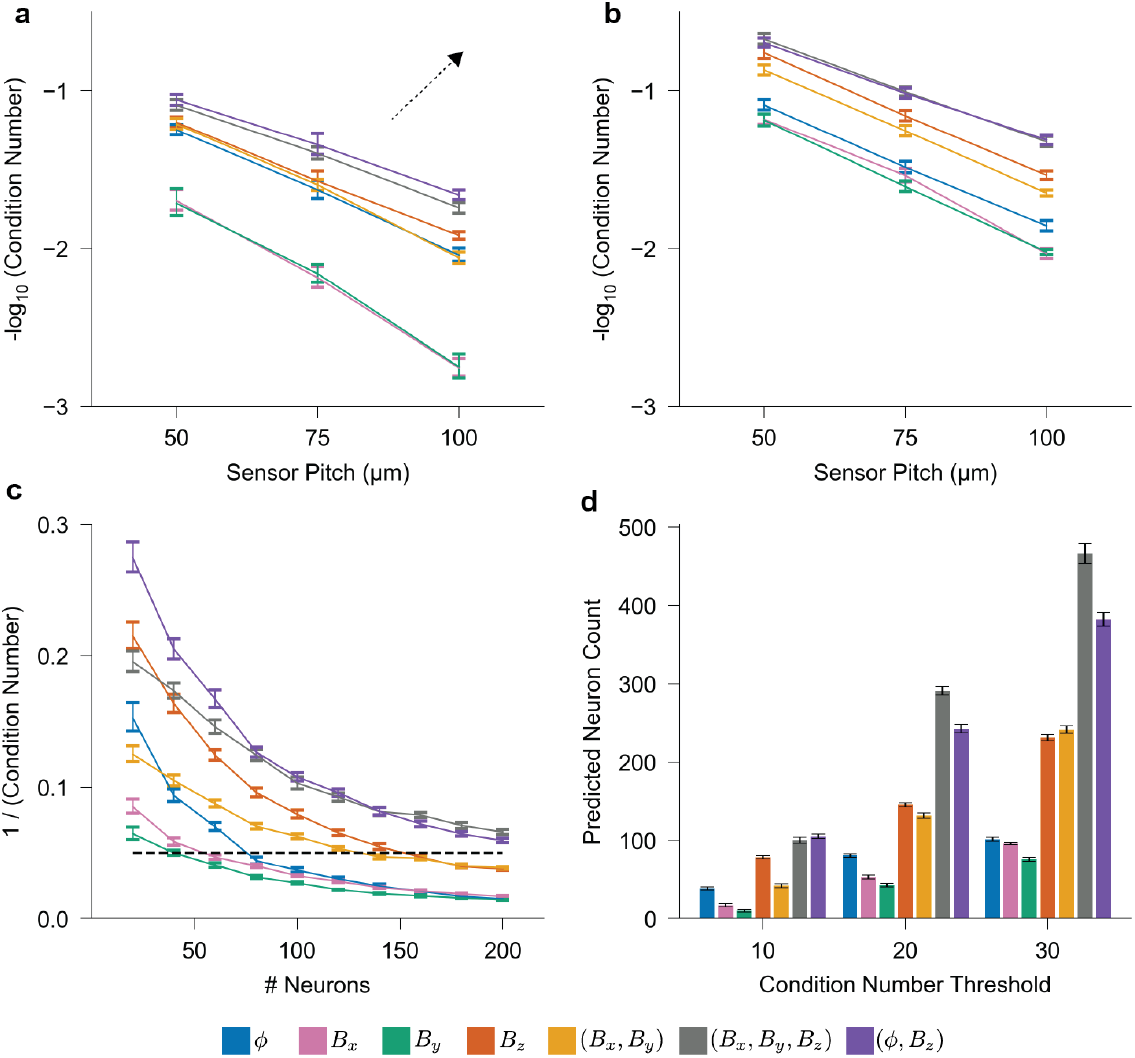
Multi-source discrimination analysis of neural electrical potential and magnetic field templates. **a**, Negative logarithm of the average condition number for channel matrices constructed from constant-density populations of *N*_*C*_ = 50, 113, and 200 cells (Cells 0–5 from SFig. 1) corresponding to sensor array pitch of 50, 75, and 100 *µ*m, respectively, for signal types *s∈ {ϕ, B*_*x*_, *B*_*y*_, *B*_*z*_, (*B*_*x*_, *B*_*y*_), (*B*_*x*_, *B*_*y*_, *B*_*z*_), (*ϕ, B*_*z*_) *}* (*N* = 30). The dashed arrow highlights the direction of a more favorable trade-off between neuronal distinguishability and coverage area. **b**, Same analysis as in **a**, but using only Cells 2 and 3 from SFig. 1. **c**, Inverse condition number as a function of cell count for arrays with 75-*µ*m pitch, using Cells 2 and 3 from SFig. 1 (*N* = 30). **d**, Maximum number of distinguishable neurons at various condition number thresholds, estimated via bootstrap resampling of the data shown in **c**.

The comparable performance of the *ϕ* and *B*_*z*_ arrays is initially counterintuitive, as our pair-wise analysis suggested a more substantial advantage for magnetic field measurements. To further investigate this discrepancy, Fig. 3b presents results using only neurons with prominent, elongated processes (Cells 2 and 3 from SFig. 1), which are more likely to generate spatially distinct magnetic field templates. In this subset, all individual magnetic components, as well as their combinations, exhibit improved trade-off performance relative to the electrical potential, which shows only minimal changes. In particular, the *B*_*z*_ array is now significantly better conditioned than the *ϕ* array across all coverage areas.

Figure 3c presents results for the second experimental scenario, where the sensor pitch is fixed at 75 *µ*m and 20 to 200 neurons are sampled from Cells 2 and 3 from SFig. 1 to assess the effect of increasing cell density. Consistent with the coverage trade-off results, the (*ϕ, B*_*z*_) and (*B*_*x*_, *B*_*y*_, *B*_*z*_) arrays yield the most favorable performance compared to the *ϕ*-only array. The (*B*_*x*_, *B*_*y*_) and *B*_*z*_ arrays also offer moderate improvements. Importantly, the relative advantage of magnetic and hybrid arrays increases with cell density, underscoring their robustness in resolving neurons in densely populated networks.

By applying a condition number threshold (for example, the dashed line in Fig. 3c) to define a minimal level of distinguishability, we estimate the maximum number of neurons that can be reliably separated for each signal type using bootstrap resampling of the data in Fig. 3c. Figure 3d summarizes these results, showing the maximum distinguishable neuron count across a range of thresholds. At nearly all tested thresholds, the *B*_*z*_ array supports approximately twice as many neurons as the *ϕ* array, while the (*ϕ, B*_*z*_) and (*B*_*x*_, *B*_*y*_, *B*_*z*_) arrays support up to three to four times more neurons. However, these multimodal arrays require two to three times as many sensors as the conventional *ϕ*-only configuration. In contrast, for cortical neuron populations, the *B*_*z*_ array outperforms the *ϕ* array even when constrained to the same number of sensors. These findings suggest that *B*_*z*_ arrays provide an efficient and scalable solution for large-scale neural recordings, particularly under practical constraints such as limited data bandwidth and sensor fabrication complexity.

### Spike Sorting

In standard electrophysiology experiments, microelectrode arrays or probes composed of *N*_*R*_ recording sites are used to capture extracellular signals from a local population of *N*_*C*_ cells. Spike sorting refers to the process of detecting, clustering, and assigning spike events to individual neurons based on similarities in their spatiotemporal patterns across the array. This approach decodes neural activity using spike timing and firing rates rather than relying solely on threshold-based event detection^45^. Building on our previous findings—namely, that dense populations of cortical neurons are more readily distinguishable via their extracellular magnetic fields than their electrical potentials, and that combining complementary electrical and magnetic recordings further enhances separability—we now investigate how these differences translate to performance in *in silico* spike sorting.

We simulate spike sorting using both *in vitro* and *in vivo* array configurations. The *in vitro* arrays are square, planar sensor arrays with varying pitches (Fig. 4a) whereas the *in vivo* array replicates the configuration of a Neuropixel 1.0 probe with 128 measurement sites^6^ (Fig. 5a). This setup allows us to compare sorting performance across different spatial layouts and experimental conditions. Simulated recordings are generated by randomly distributing a population of neurons (Cells 2 and 3 from SFig. 1) across the arrays, summing and concatenating their Poisson-generated spikes, and subsequently adding noise. Spike sorting is performed using Kilosort2, a high-performance algorithm based on template matching that effectively resolves overlapping spikes^46^. Performance is evaluated using two metrics: overall sorting accuracy, defined as the match rate between predicted and ground truth spike trains, and the number of well-detected cells, defined as units with match rates exceeding 80%^47^. These well-detected cells, which typically correspond to one-to-one matches with ground truth neurons, reduce the need for manual curation, such as merging or splitting spike templates^48^.

**Figure 4.**
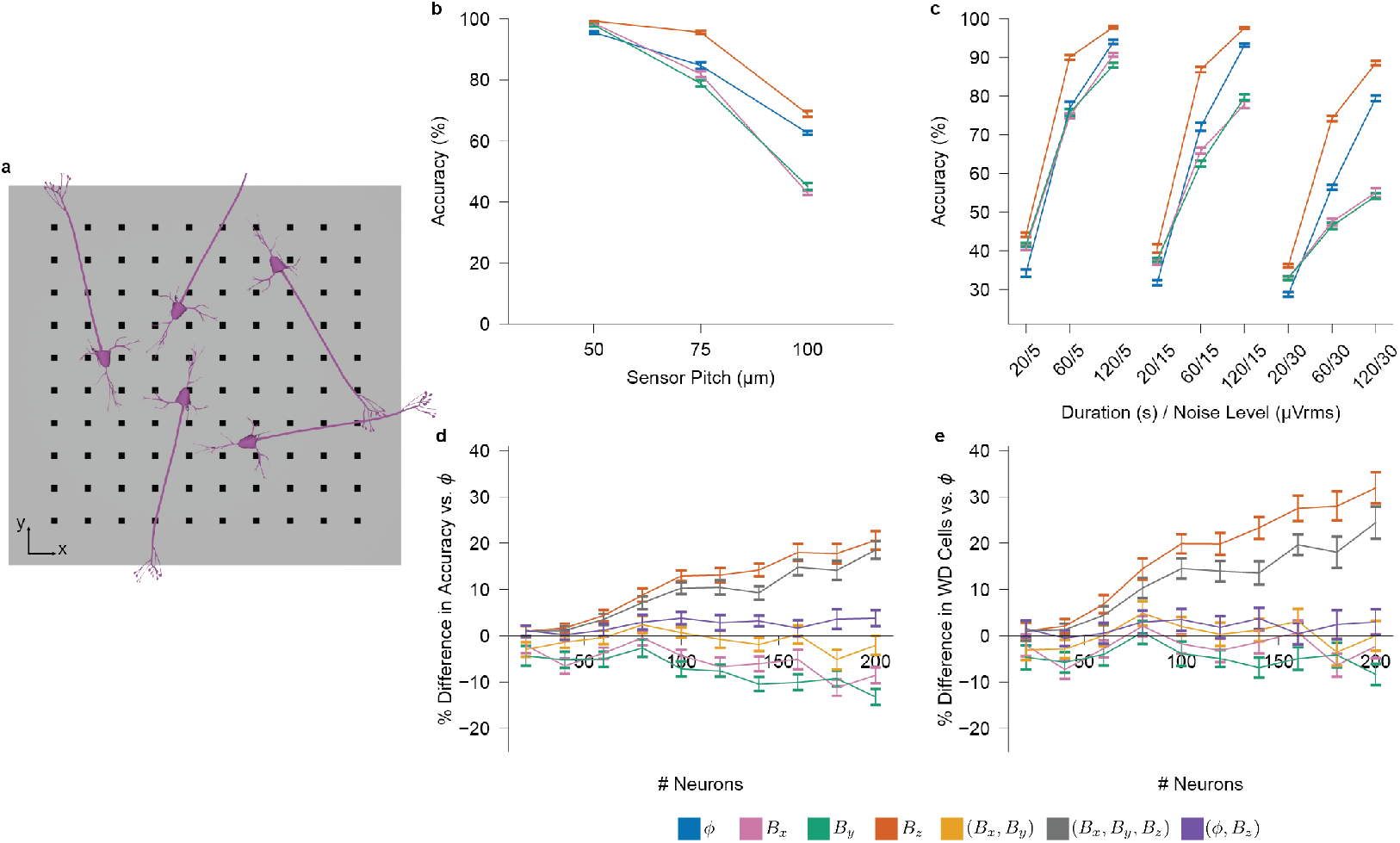
Simulated spike sorting for *in vitro* neural probes. **a**, Simulation setup for planar *in vitro* arrays. The array consists of a 10 *×* 10 grid of measurement points, with neurons randomly positioned and oriented above the grid. **b**, Spike sorting accuracy for *s ∈ {ϕ, B*_*x*_, *B*_*y*_, *B*_*z*_*}* as a function of sensor pitch (50, 75, and 100 *µ*m; corresponding to 50, 113, 200 cells at constant density; 60 s duration, 15 *µ*V_rms_ noise level). The noise level for magnetic templates is in units of tesla and is normalized such that their average SNR matches that of the *ϕ* template. **c**, Sorting accuracy as a function of recording duration (20, 60, 120 s) and noise level (5, 15, 30 *µ*V_rms_) for each signal type (200 cells, 75 *µ*m pitch). **d**, Percent difference in spike sorting accuracy relative to *ϕ* for signal types *s ∈ {B*_*x*_, *B*_*y*_, *B*_*z*_, (*B*_*x*_, *B*_*y*_), (*B*_*x*_, *B*_*y*_, *B*_*z*_), (*ϕ, B*_*z*_) *}* as a function of cell count (75 *µ*m pitch, 60 s duration, 15 *µ*V_rms_ noise level). **e**, Percent difference in the number of well-detected cells (match rate *>* 80%) under the same conditions as in **d**. *N* = 10 for all panels.

**Figure 5.**
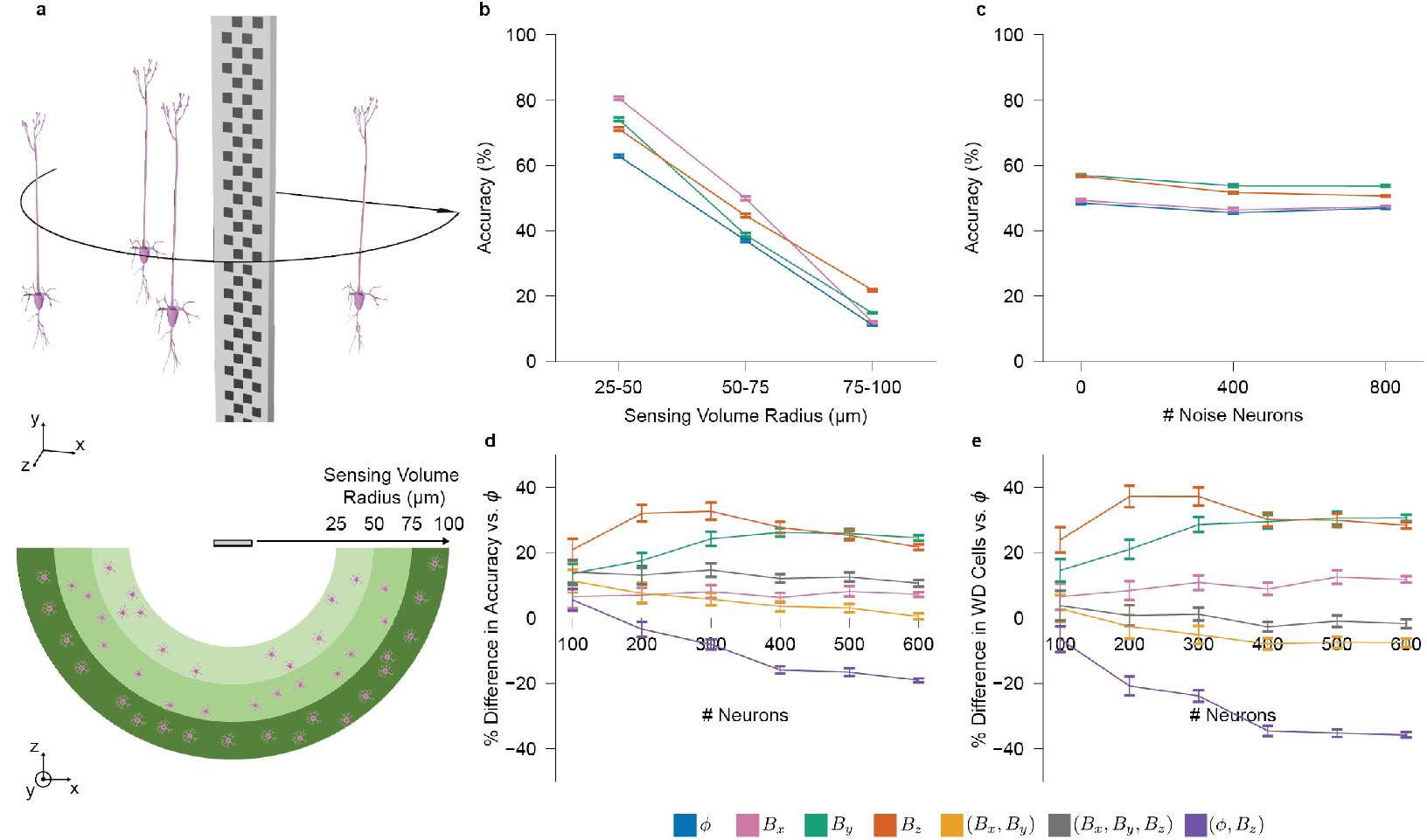
Simulated spike sorting for *in vivo* neural probes. **a**, Simulation setup for a Neuropixel-style *in vivo* probe. Neurons are distributed within a defined radius on the front side of the probe. **b**, Spike sorting accuracy as a function of distance from the probe for *s∈ {ϕ, B*_*x*_, *B*_*y*_, *B*_*z*_*}* (400 cells, 120 s duration, 15 *µ*V_rms_ noise level). **c**, Sorting accuracy as a function of noise neuron count (neurons located beyond 100 *µ*m from the probe) for each signal type (400 signal neurons within 10-100 *µ*m radius, 120 s duration). **d**, Percent difference in sorting accuracy relative to *ϕ* for signal types *s ∈ {B*_*x*_, *B*_*y*_, *B*_*z*_, (*B*_*x*_, *B*_*y*_), (*B*_*x*_, *B*_*y*_, *B*_*z*_), (*ϕ, B*_*z*_) *}*, as a function of cell count (120 s duration, 15 *µ*V_rms_ noise level, 10–100 *µ*m cell radius). **e**, Percent difference in the number of well-detected cells under the same conditions as **d**. *N* = 10 for all panels.

Figure 4b presents the trade-off between spike sorting accuracy and coverage area for *in vitro* arrays with a fixed number of sensors. Across all coverage areas, the normal magnetic field *B*_*z*_ consistently yields higher sorting accuracy than the electrical potential *ϕ*, while the planar magnetic components *B*_*x*_ and *B*_*y*_ perform worse than *ϕ* except at the smallest coverage area. To further examine these trends, we evaluate how recording duration and noise level affect sorting performance. As shown in Fig. 4c, the relative ranking of spike sorting accuracy for all signal types remains consistent as a function of these variations. As expected, accuracy improves with longer recording durations and degrades with increased noise across all signal types. While the performance of *ϕ* and *B*_*z*_ declines gradually with increasing noise, the degradation in *B*_*x*_ and *B*_*y*_ is significantly steeper, reflecting the same pronounced sensitivity to coverage area (Fig. 4b).

These findings align with our earlier multi-source discrimination analysis, in which *B*_*x*_ and *B*_*y*_ exhibited significantly larger condition numbers—indicating poor signal separability—while the normal magnetic field *B*_*z*_ is consistently better conditioned. Although noise was not explicitly considered in that analysis, the sharper decline in sorting accuracy for *B*_*x*_ and *B*_*y*_ under noise reinforces their ill-conditioned nature. This demonstrates that poorly conditioned channels are more susceptible to noise while their performance can be substantially improved with sufficiently small sensor pitch. Overall, these findings support the condition number as a strong predictor of spike sorting algorithms’ ability to reliability separate and identify neural signals in practical settings.

Figure 4d evaluates how increasing cell density within a fixed coverage area affects sorting accuracy for *in vitro* arrays. To facilitate comparison, we plot the percentage difference in spike sorting accuracy between each signal type and the *ϕ* array. At low cell densities, the *B*_*z*_ and (*B*_*x*_, *B*_*y*_, *B*_*z*_) arrays perform comparably to the *ϕ* array; however, at the highest evaluated density, they yield nearly a 20% increase in sorting accuracy. In contrast, the *B*_*x*_ and *B*_*y*_ arrays consistently underperform, while the (*B*_*x*_, *B*_*y*_) and (*ϕ, B*_*z*_) arrays remain comparable to the *ϕ* array. A similar trend appears in Fig. 4e, where the *B*_*z*_ and (*B*_*x*_, *B*_*y*_, *B*_*z*_) arrays yield 20–30% more well-detected cells than the *ϕ* array, while other multimodal arrays show marginal or no improvement.

These results reaffirm the effectiveness of measuring the normal magnetic field component *B*_*z*_, alone or in combination with other fields, which is consistent with our distinguishability analysis identifying *B*_*z*_ as best suited for recording from dense neural networks. However, multimodal arrays—particularly (*ϕ, B*_*z*_)—underperform relative to predictions from the distinguishability analysis. This discrepancy likely reflects limitations of the spike sorting algorithm rather than the signal content itself. Since Kilosort incorporates sensor position into its model, it may struggle with co-located heterogeneous signals. Additionally, the two- to three-fold increase in sensor count for multimodal arrays may further challenge algorithmic performance.

Next, we turn to *in vivo* settings, where probes sample from a three-dimensional (3D) volume (Fig. 5a, top), and only neurons near the probe are reliably detected, while distant cells contribute background noise^49^. Because sorting accuracy depends on the number of detectable neurons, we define an effective sensing radius—the distance beyond which neurons are excluded and treated as noise—illustrated in the spatial setup of Fig. 5a (bottom), where accuracy is evaluated across radial distances. Fig. 5b shows that sorting accuracy for individual components (*ϕ, B*_*x*_, *B*_*y*_, *B*_*z*_) decreases linearly with radial distance, dropping below 25% between 75 and 100 *µ*m. We therefore define 100 *µ*m as the effective sensing radius, using 25% as a conservative threshold below which spike assignments are considered unreliable. To determine whether neurons beyond this radius should be treated as noise, we vary their number beyond 100 *µ*m (Fig. 5c). Sorting accuracy remain largely unaffected across signal types, supporting the exclusion of biological noise from further simulations

Unlike the *in vitro* case, where we examine trade-offs between sorting accuracy and both coverage and cell density, the *in vivo* analysis focuses solely on the trade-off with cell density, as coverage is fixed by the probe geometry. As shown in Figure 5d, individual magnetic field arrays achieve up to 25–35% higher sorting accuracy than the *ϕ* array, even at the highest evaluated cell densities. In contrast, multimodal arrays underperform relative to the individual components, likely due to the same algorithmic limitations observed *in vitro*. While the (*B*_*x*_, *B*_*y*_, *B*_*z*_) array provide a modest 10% improvement over *ϕ*, the (*ϕ, B*_*z*_) array performs significantly worse. Figure 5e shows that well-detected cell counts follow similar trends.

In contrast to the *in vitro* setting, while the normal magnetic field *B*_*z*_ continues to outperform *ϕ*, it is no longer a clear outlier relative to the planar components. This shift likely stems from the three-dimensional cellular arrangement *in vivo*, where a neuron’s magnetic field may be oriented either normal or tangential to the probe surface—unlike the *in vitro* setting, where neurons predominantly lie in a plane orthogonal to the *B*_*z*_ sensing axis.

Together, these results demonstrate that the theoretical advantages of magnetic field sensing for neuronal discrimination extend to practical spike sorting scenarios, even when using algorithms originally developed for electrical potential recordings. Across both *in vitro* and *in vivo* settings, the normal magnetic field component *B*_*z*_ consistently enhances sorting accuracy and increases the number of reliably identified neurons, particularly in dense neural populations. While performance gains are partially limited by algorithmic assumptions–especially in multimodal, co-located configurations—the robustness of *B*_*z*_ across spatial scales and noise conditions highlights its value for next-generation neural interfaces. Future improvements may be realized through dedicated algorithmic development tailored to the unique characteristics of vectorial and composite magnetic recordings.

### Morphological Reconstruction

Thus far, we have examined how fundamental differences between neural electrical potentials and magnetic fields affect the separability of individual neurons within large populations. We now shift focus to the single-cell level, exploring how the distinctive properties of neural magnetic fields can also inform the structural characterization of individual neurons. In particular, we explore morphology reconstruction, the task of estimating the spatial configuration of a neuron and its major processes. Traditionally, this task requires dense electrode arrays with inter-electrode spacing below 20 *µ*m, along with complex algorithms or antidromic stimulation, due to the small radius and weak extracellular potential signal of neuronal processes relative to the soma^50–52^. Here, we investigate whether magnetic field measurements can facilitate morphology reconstruction using sparser arrays, which are more compatible with large-scale network recordings.

As shown in Fig. 1b, planar *in vitro* arrays measuring normal magnetic fields show opposite polarities on either side of a major process, allowing its localization by detecting the boundary between positive and negative versions of the extracellular spike waveform. To estimate this boundary, we repurpose support vector machines (SVMs) for single-shot estimation—deviating from their conventional use in classification—and refer to this approach as General Neural Boundary Estimation (GNBE). SVMs are well-suited due to their off-the-shelf availability, training-free use in this context, and ability to compute contiguous, non-linear hyperplanes that better match the geometry of realistic neural processes compared to alternatives such as logistic regression or random forests^53^. In GNBE, a noisy *B*_*z*_ spike template—representing a single neuron’s estimated signal after spiking sorting—is input into the SVM. Using sensor locations and signal polarities, the SVM computes a boundary curve interpreted as the dominant process’s orientation. This estimated boundary is then compared to the ground-truth morphology, as illustrated in Fig. 6a. To assess feasibility, we simulate neurons on planar sensor arrays across varying sensor pitches and signal-to-noise ratios (SNRs), reflecting different levels of template quality.

**Figure 6.**
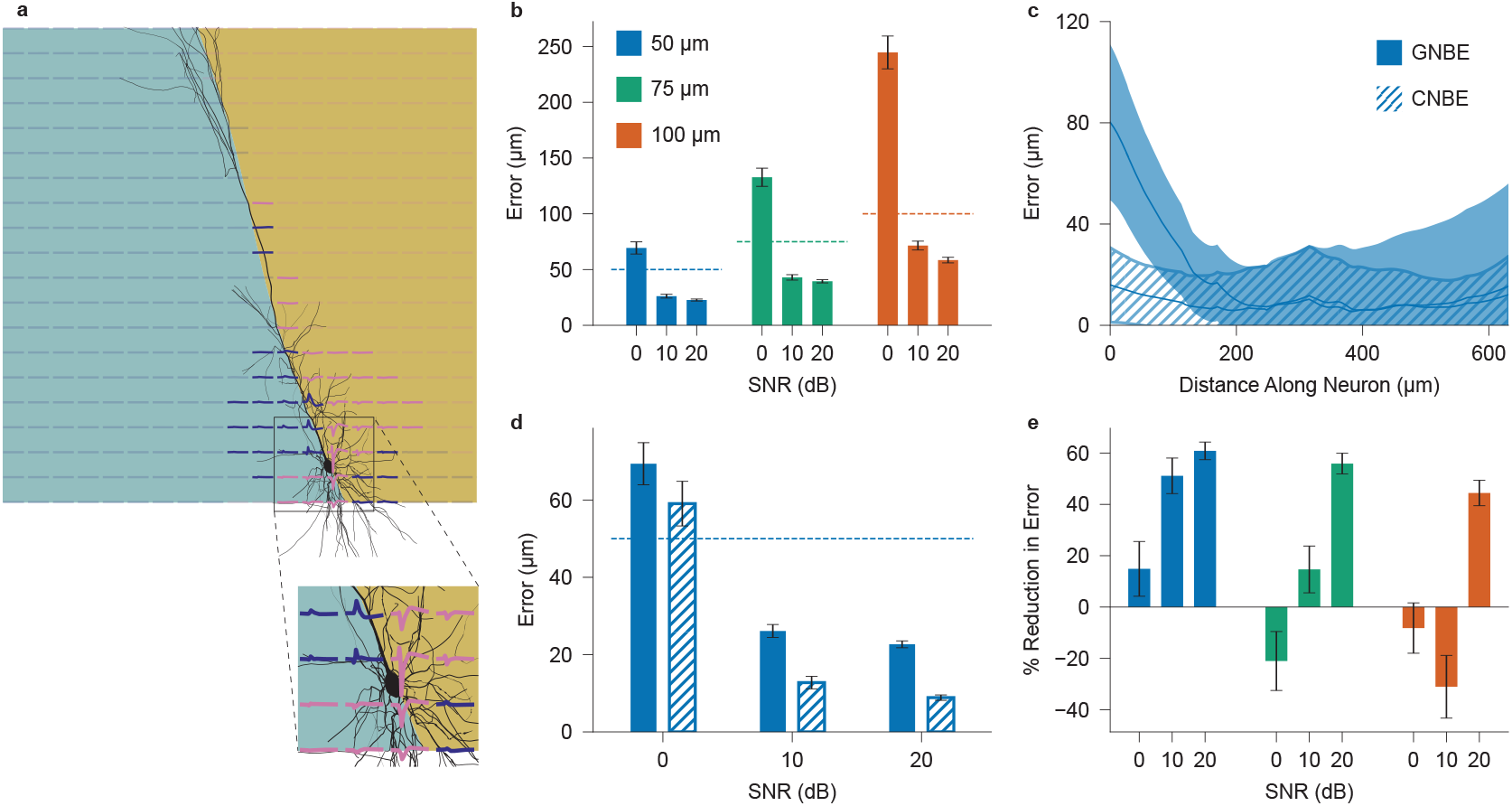
Morphology reconstruction for simulated *B*_*z*_ *in vitro* sensor arrays. **a**, Example of a decision boundary computed using the GNBE method, overlaid on the ground-truth neuron morphology. Spike signals at each sensor are shown, with positive signals in indigo and negative in pink. Inset shows how magnetic field polarities differ on opposite sides of the soma (top versus bottom) due to opposing longitudinal current flows. **b**, Reconstruction error using the GNBE algorithm for *B*_*z*_ templates with 0, 10, and 20 dB average SNR, across 20 *×* 20 sensor arrays with 50, 75, and 100 *µ*m pitch. Dashed lines indicate corresponding values of sensor pitch. **c**, Average reconstruction error as a function of distance from the soma along the apical dendrite, comparing GNBE and CNBE methods. **d**, Reconstruction error versus SNR for a 20 *×* 20 *B*_*z*_ array with 50 *µ*m pitch, comparing GNBE and CNBE methods. **e**, Percent reduction in reconstruction error achieved by CNBE method under the same conditions as in **a**. *N* = 300 randomly oriented instances of Cell 2 from SFig. 1. Standard error is plotted for **b, d, e**, and standard deviation is shown for **c**.

Figure 6b shows the reconstruction error, defined as the average distance between each neuronal segment and the predicted boundary curve, as a function of template SNR and sensor pitch. For moderate (10 dB) and high (20 dB) SNRs, the reconstruction error is smaller than the sensor pitch for all pitch values tested, demonstrating super-resolution accuracy. At low SNR (0 dB), performance degrades considerably but improves with decreasing senor pitch. These results confirm that normal magnetic field sensing is highly effective for morphology reconstruction, enabling accurate estimation of neuronal morphology even with sparse and noisy templates using off-the-shelf algorithms.

Off-the-shelf GNBE-based reconstructions exhibit consistent inaccuracies near the soma, due to dendrites connected to the soma and oriented away from the primary neural process generating magnetic fields that rotate in the opposite direction (Fig. 6a inset). To address this, we introduce a second-pass algorithm, referred to as Customized Neural Boundary Estimation (CNBE), that leverages known properties of cortical magnetic fields. CNBE proceeds in four steps: apply GNBE to obtain an initial boundary estimate; use this estimate to approximate the soma location and the direction of the major process; discard sensors on the side opposite the major process—typically associated with dendritic fields (Fig. 6a, insert); and re-estimate the boundary using only the remaining sensors. Full algorithmic details are provided in the Methods section.

Figure 6c shows reconstruction error along the major process as a function of distance from the soma, comparing GNBE and CNBE algorithms for a 50 *µ*m-pitch array. CNBE effectively eliminates the elevated error near the soma observed with the naive method. Figure 6d summarizes reconstruction error across all SNR levels for both algorithms. With CNBE, the error is reduced to approximately 20% of the pitch at moderate and high SNR, with modest improvement even at low SNR. Figure 6e quantifies the percentage reduction in reconstruction error achieved by the CNBE algorithm across all sensor pitches.

These results demonstrate that CNBE yields significant improvement when applied to dense arrays or high-SNR templates. However, its performance may degrade on sparse arrays or low-SNR templates, as it depends on the initial GNBE estimate being reasonably accurate. When effective, CNBE reduces reconstruction error by over 50%, highlighting the benefit of customized over off-the-shelf algorithms. Overall, the findings reinforce the potential of normal magnetic field sensing for morphological reconstruction, owing to its origin in longitudinal—rather than transmembrane—currents.

## DISCUSSION

Our theoretical analysis reveals that extracellular magnetic fields and electrical potentials originate from distinct components of the intracellular surface current density—specifically, longitudinal and transmembrane currents, respectively. This biophysical distinction, derived via Green’s theorem, suggests that magnetic fields are particularly sensitive to extended neuronal structures such as dendrites and axons, whereas electrical potentials are dominated by somatic activity. As a result, magnetic recordings, especially from the normal component of the field, can provide richer spatial and morphological information for certain cell types. The rotational nature of magnetic fields also leads to more spatially diverse signal patterns, which may aid in resolving overlapping sources within densely packed neuronal populations. These insights motivate the integration of electrical and magnetic recordings to enhance both the resolution and interpretability of extracellular measurements.

To further characterize these differences, we examined how electrical potentials and magnetic fields scale with distance from their sources. While the electric field from transmembrane currents decays rapidly due to its quadrupole-like structure, the magnetic field—arising from longitudinal currents—exhibits dipole-like behavior with a slower decay. Simulations using realistic morphologies showed that the normal magnetic field decays roughly as 1*/R*^2^, compared to 1*/R*^2.5^ for the electrical potential. This difference in spatial reach suggests that the normal magnetic field may retain more information at greater distances, offering advantages for recording in larger volumes and sparser sensor configurations.

Building on this foundation, we quantify how these signal properties influence the ability to distinguish individual neurons. We evaluate pairwise neuronal resolution using a spatial similarity metric applied to spike templates from morphologically realistic cells. While electrical potentials produce highly symmetric and correlated templates due to their somatic origin, magnetic fields— particularly, the normal magnetic field—capture more complex spatial features. This results in significantly improved resolution, with the normal magnetic field reducing the minimum resolvable pairwise separation distance by up to nine-fold compared to the electrical potential. Planar components also improve resolution but are more sensitive to neuronal orientation. These results underscore the fundamental advantage of measuring normal magnetic fields for distinguishing neurons in dense networks and establish a strong basis for extending the analysis to larger populations.

To assess cell distinguishability at the network level, we apply the condition number—a communication theory metric for quantifying signal separability—to simulated spike templates of cortical populations. Across varying array coverages and cell densities, normal magnetic field templates consistently outperform electrical potentials, especially in populations with elongated neuronal processes. Planar magnetic components perform worse due to orientation-dependent blind spots, where signals align orthogonally to the sensor axes. Although the advantage of normal magnetic sensing diminishes in populations lacking strong longitudinal structures, combining magnetic and electrical signals reliably improves separability, underscoring their complementary strengths. While multimodal configurations yield the highest gains, they require substantially more sensors. Notably, sensing the normal magnetic field alone provides a strong trade-off, exceeding the performance of electrical potentials even under matched hardware constraints. These results underscore the practical utility of magnetic recordings for dense neural monitoring and support their scalability in settings limited by data throughput and fabrication complexity

We next evaluate whether the theoretical advantages of magnetic field sensing translate to practical spike sorting performance using simulated recordings processed with Kilosort2. Across both *in vitro* and *in vivo* configurations, recordings of the normal magnetic field consistently improve spike sorting accuracy and increase the number of well-detected units, especially in dense neuronal populations. Planar magnetic components perform less reliably in the *in vitro* setting but match the performance of the normal magnetic field *in vivo*, likely due to the three-dimensional organization of neural tissue, smaller sensor pitch, and greater sensitivity to distant sources. Multimodal recordings that combine electrical potential and magnetic field signals fall short of theoretical expectations, likely due to current algorithmic limitations in handling co-located, heterogeneous inputs. Nonetheless, spike sorting based on the normal magnetic field remains robust to noise, recording duration, and population density, suggesting that targeted algorithmic improvements could further enhance its utility.

Beyond enhancing conventional neural analyses, we explore how the unique characteristics of magnetic field measurements—particularly the solenoidal nature of the normal component—can support structural inference at the single-cell level. Because this component exhibits polarity reversals across extended neural processes, we leverage this spatial pattern to estimate the orientation of major morphological structures. Using support vector machines (SVMs) repurposed for polarity-based boundary estimation, we achieve super-resolution accuracy even with low-SNR spike templates. A second-pass refinement, which excludes signals likely associated with dendritic polarity inversion near the soma, further improves performance. Under favorable conditions, this approach reduces reconstruction error by more than 50%. These results suggest that recordings of the normal magnetic field, even from sparse sensor arrays, can enable coarse morphology estimation in *in vitro* settings and offer structural insights that complement traditional spike-based analyses.

Our results point to a critical and underexplored direction for future sensor development: the need for highly sensitive magnetic sensors for the normal component of neural magnetic fields. Currently, no magnetic sensor technology offers the necessary combination of sensitivity, spatial resolution, and low noise to enable single-shot recordings of cortical or peripheral activity. Magnetoresistive (MR) sensors are among the most promising candidates due to their miniature size, scalability to large arrays, and compatibility with CMOS fabrication^20,22^. However, most MR devices are optimized for planar field detection. Normal-field MR sensors remain far less explored and developed and, where fabricated, typically exhibit reduced sensitivity^54^. Our findings demonstrate that normal magnetic field recordings provide substantial advantages in neuronal separability, spatial reach, and morphology reconstruction—benefits that go well beyond their known clinical value. These insights make a compelling case for renewed focus on the design and optimization of normal-field MR sensors, which could greatly expand the capabilities of implanted neural recording technologies.

## Supporting information

Supplemental Proofs and Figure 1

## RESOURCE AVAILABILITY

### Lead contact

Requests for further information and resources should be directed to and will be fulfilled by the lead contacts, Ziad Ali and Ada Poon.

### Data and code availability

Any information required to reanalyze the data reported in this paper is available from the lead contacts upon request.

## ACKNOWLEDGMENTS

This work was funded by the Knight-Hennessy Scholarship, the NSF GRFP award, and a seed grant from the Wu Tsai Neurosciences Institute. The authors thank all members of the lab for their support.

## AUTHOR CONTRIBUTIONS

Conceptualization, Z.A and A.S.Y.P; methodology, Z.A and A.S.Y.P; investigation and modeling, Z.A and A.S.Y.P; writing, Z.A and A.S.Y.P; funding acquisition, Z.A and A.S.Y.P; supervision, A.S.Y.P.

## DECLARATION OF GENERATIVE AI AND AI-ASSISTED TECHNOLOGIES

During the course of preparing this manuscript, the author(s) used OpenAI’s ChatGPT for the purpose of editing suggestions and checking grammar. Following the use of this tool/service, the author(s) formally reviewed the content for its accuracy and edited it as necessary. The author(s) take full responsibility for all the content of this publication.

## SUPPLEMENTAL INFORMATION INDEX

Figure S1. Reference for the different cell types used in the computational studies.

## STAR METHODS

### Neuron Simulation and Extracellular Calculations

To calculate neural action potentials we used the NEURON simulator library^37^, and to calculate extracellular electrical potentials we used the line-source approximation^55^ within the LFPy module^38^. Spikes were elicited by injecting current into the soma using NEURON’s IClamp point process. We used the MEArec package^56^ to generate simulated extracellular electrical potential recordings from large populations and modified it to calculate and incorporate extracellular magnetic field data. Magnetic field calculations were made using the Biot-Savart Law approximation^57^ updated for a neuron model as shown in Eqn. 18, which describes the magnetic field at an extracellular coordinate **r** in terms of the neuron compartmental axial current *I*_*a*_, direction vector **d**, and relative position vector **r**^*′*^ (pointing from the compartment to the calculation point) for each compartment *n*.

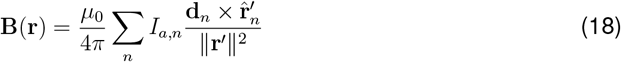

For both *in vitro* and *in vivo* arrays, cell morphologies were modified so as to neither intersect the plane of the array nor extend, unrealistically, upwards into space. For *in vitro* arrays, all cells were flattened to within a z-dimension of 5–25 *µ*m above the plane of the sensors. For the *in vivo* arrays, all cellular processes that would have intersected the shank were made to instead wrap around the nearest shank boundary. Cells were also flattened so as to not cross within 5 *µ*m of the front-face of the shank.

### Similarity Calculation and Template Creation

Simulated cells were suspended 15 *µ*m (soma as the origin) over a 1,500 1,500 grid of measurement points with 2 *µ*m pitch and flattened. This setup was intended to model a traditional *in vitro* electrophysiology experiment but with near-infinitesimal resolution. The signal traces of each sensor for the spike duration (224 timesteps, *dt* = 31.25 *µ*s, 7 ms total) were combined to form a spike template 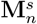 for each neuron *n* and signal type *s* (2,250,000*×*224, *n* = 0 to 5, *s ∈ {ϕ, B*_*x*_, *B*_*y*_, *B*_*z*_*}*). To create 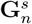, we defined the *T*_*sweep*_ transformation over *θ* from 0 to 360^*°*^ in 15^*°*^ increments. Translation coordinates Δ**r** for *T*_*trans*_(Δ**r**) were defined over a coarse grid from ( *−*400 *µ*m, *−*400 *µ*m) to (400 *µ*m, 400 *µ*m) with 50 *µ*m pitch and a fine grid from ( *−*200 *µ*m, *−* 200 *µ*m) to (200 *µ*m, 200 *µ*m) with 20 *µ*m pitch (in the *x*-*y* plane).

In generating templates for rotated cells, we had to contend with the fact that rotated sensor coordinates no longer necessarily aligned with the original sensor coordinates. Therefore, after obtaining the rotated coordinates, the *K*-nearest rotated sensors (*K* = 5) of each original sensor were determined. Then, the new template value *p*_0,*θ*_ at each original sensor location was calculated as the average of the data at the *K*-nearest neighbors inversely weighted by distance as described in (19) (for data *p*_*k*_ at the *K*-nearest neighbors with distance *d*_*k*_ and weights *w*_*k*_).

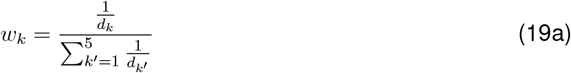

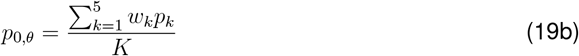

### Spike Sorting Recording Generation

To generate the simulated recording data for spike sorting, we first created template libraries in which a sensor configuration was chosen (such as a 10 *×* 10 planar grid of sensors with 50, 75, or 100 *µ*m pitch, or a Neuropixel 1.0 probe), a cell type was specified (Cell 2 or 3 from SFig. 1), and then *n* different spike templates of signal *s*—for *s ∈ {ϕ, B*_*x*_, *B*_*y*_, *B*_*z*_*}* —for that cell were generated by randomly varying the location and rotation angle of the cell. For the Neuropixel, cells were placed only within some radius in front of the probe (volume defined by a half-cylinder) due to the inability of our simulation environment to adequately model transmembrane currents originating behind the probe (which would theoretically travel around it to reach microelectrodes rather than directly through it). All cells were flattened and/or wrapped around the sensor array.

Following the generation of a template library for each cell for a specified sensor configuration, *N*_*C*_ cell templates were randomly sampled across the combined templates of all selected cell types to represent the *N*_*C*_ neurons being simultaneously measured by the probe. A Poisson spike train was generated using MEArec for each template with a rate chosen from a normal distribution (mean = 5 Hz, standard deviation = 1 Hz) and the resultant extracellular signals were then combined to produce a recording. Spikes violating a set refractory period of 2 ms were removed.

Gaussian thermal noise was generated and added to voltage recordings according to a set noise power value. The range of noise powers used was chosen based on fundamental thermal noise properties of resistors. A typical impedance value (at 1 kHz) for an electrode with a size on the order of 10 *µ*m is generally between 100 kOhm to over 1 MOhm depending on the electrode material^6,58,59^. The power spectral density of thermal noise from a resistor is 4*kTR*, so for an electrode measuring a signal in a 32 kHz bandwidth (equivalent to our *dt* for spike generation; data was band-pass filtered during spike sorting), this impedance range approximately corresponds to a noise RMS voltage range of 7.35 to 32.9 *µ*V_rms_. We chose to simulate from 5 to 30 *µ*V_rms_. While realistic electrode impedances scale as a function of frequency, assuming a constant resistance provides us with a simple and conservative estimate. Notably, the Neuropixel 1.0 probe reports approximately 7.1 *µ*V_rms_ noise for an *in vivo* implant within a 300 Hz to 10 kHz bandwidth^60^—this corresponds to 12.9 *µ*V_rms_ when the bandwidth is changed to 32 kHz. We therefore chose 15 *µ*V_rms_ as our default noise level for the *in vitro* and *in vivo* simulations.

For magnetic sensor noise, to make a fair comparison considering the difference in units— volts vs. tesla—the average signal power across the sensors for the entire recording duration was calculated for each magnetic field component (*B*_*x*_, *B*_*y*_, *B*_*z*_) and the *ϕ* data. The magnitude of the magnetic field signals was then scaled up so that the average signal power of the magnetic field data matched the average signal power of the voltage data (numerically, ignoring units). After the additive sensor noise was included, each field had an equivalent average SNR. This also ensured that the scale of the signals was approximately identical when fed into the sorting algorithm which was important to minimize unwanted variance. For multimodal measurements, the fields were scaled according to the signal type with the largest average power (so as to preserve the relative magnitudes of each field type).

### Simulated Spike Sorting

Kilosort2^46^ was used within the SpikeInterface^47^ framework for spike sorting. The only significant modification we made to the Kilosort algorithm was with regards to spike detection based on threshold crossing: whereas the original algorithm detected only negative spikes, we set it to detect either negative or positive spikes for magnetic fields. Implementing this change for voltage signals worsened the overall performance (the largest spikes, usually near the soma, typically have negative polarity), and so was not included, but it was essential for magnetic fields given that their largest spikes can have different polarities depending on the direction of longitudinal currents relative to the sensor positions.

We performed a grid search across Kilosort2’s detection threshold and minimum spiking frequency variables for each combination of recording parameters—recording duration, number of neurons, noise level, and sensor configuration—to attempt to find the close-to-optimal sorting performance for each parameter set. We randomly generated *N* = 10 recordings for each combination of parameters and then, for each recording, determined the average sorting accuracy, defined as true positive / (true positive + false positive + false negative) for predicted spikes, and the number of well-detected units, defined as having greater than 80% match between their ground-truth spike train and spike-sorted spike train^47^. The highest-accuracy sorts and highest well-detected-count sorts for each signal *s ∈ {ϕ, B*_*x*_, *B*_*y*_, *B*_*z*_*}* across the grid search were isolated and compared to get an estimate of the upper bound of sorting performance per field type, regardless of sorting condition.

### Morphology Reconstruction Modified Algorithm

The CNBE algorithm to enhance the performance of the GNBE approach for morphology reconstruction proceeds as follows. First, a naive SVM boundary is calculated. Next, the location of the axon hillock (the region of the cell typically responsible for the strongest magnetic field signal^61^) is estimated by finding the weighted center of the coordinates of the 4 sensors that measured the signals with the greatest average power. This procedure is repeated for the next 96 highest power sensors, producing 24 points that were estimated to lie close to, or in the general direction of, the neuron’s actual morphology. The 24 points along the naive SVM boundary lying closest to these points are then determined, and “outliers”—with an average distance to their 4 nearest neighbors greater than 50 *µ*m—are removed. A line of best fit that runs through the remainder of these 24 points and the axon hillock coordinate is then calculated (strongly weighted to pass close to the axon hillock). A line perpendicular to this best fit line and lying 10 *µ*m “behind” the axon hillock (the opposite direction of the vector pointing from the axon hillock to the 24 points) is then constructed—any sensor behind this line is then excluded from a second SVM calculation, which produces an updated morphology estimate.

## References

1. Hochberg, L.R., Serruya, M.D., Friehs, G.M., Mukand, J.A., Saleh, M., Caplan, A.H., Branner, A., Chen, D., Penn, R.D., and Donoghue, J.P. (2006). Neuronal ensemble control of prosthetic devices by a human with tetraplegia. Nature 442, 164–171. URL: https://doi.org/10.1038/nature04970. doi: 10.1038/nature04970.

2. Willett, F.R., Kunz, E.M., Fan, C., Avansino, D.T., Wilson, G.H., Choi, E.Y., Kamdar, F., Glasser, M.F., Hochberg, L.R., Druckmann, S., Shenoy, K.V., and Henderson, J.M. (2023). A high-performance speech neuroprosthesis. Nature 620, 1031–1036. URL: https://doi.org/10.1038/s41586-023-06377-x. doi: 10.1038/s41586-023-06377-x.

3. Jepson, L., Hottowy, P., Weiner, G., Dabrowski, W., Litke, A., and Chichilnisky, E.J. (2014). High-fidelity reproduction of spatiotemporal visual signals for retinal prosthesis. Neuron 83, 87–92. URL: https://doi.org/10.1016/j.neuron.2014.04.044. doi: 10.1016/j.neuron.2014.04.044.

4. Pasley, B.N., David, S.V., Mesgarani, N., Flinker, A., Shamma, S.A., Crone, N.E., Knight, R.T., and Chang, E.F. (2012). Reconstructing speech from human auditory cortex. PLoS biology 10, e1001251.

5. Jones, K.E., Campbell, P.K., and Normann, R.A. (1992). A glass/silicon composite intra-cortical electrode array. Annals of Biomedical Engineering 20, 423–437. URL: https://doi.org/10.1007/BF02368134. doi: 10.1007/BF02368134.

6. Jun, J.J., Steinmetz, N.A., Siegle, J.H., Denman, D.J., Bauza, M., Barbarits, B., Lee, A.K., Anastassiou, C.A., Andrei, A., Aydin, Ç., Barbic, M., Blanche, T.J., Bonin, V., Couto, J., Dutta, B., Gratiy, S.L., Gutnisky, D.A., Häusser, M., Karsh, B., Ledochowitsch, P., Lopez, C.M., Mitelut, C., Musa, S., Okun, M., Pachitariu, M., Putzeys, J., Rich, P.D., Rossant, C., Sun, W.l., Svoboda, K., Carandini, M., Harris, K.D., Koch, C., O’Keefe, J., and Harris, T.D. (2017). Fully integrated silicon probes for high-density recording of neural activity. Nature 551, 232–236. URL: https://doi.org/10.1038/nature24636. doi: 10.1038/nature24636.

7. Szarowski, D., Andersen, M., Retterer, S., Spence, A., Isaacson, M., Craighead, H., Turner, J., and Shain, W. (2003). Brain responses to micro-machined silicon devices. Brain Research 983, 23–35. URL: https://www.sciencedirect.com/science/article/pii/S0006899303030233. doi: 10.1016/S0006-8993(03)03023-3.

8. Chestek, C.A., Gilja, V., Nuyujukian, P., Foster, J.D., Fan, J.M., Kaufman, M.T., Churchland, M.M., Rivera-Alvidrez, Z., Cunningham, J.P., Ryu, S.I., and Shenoy, K.V. (2011). Long-term stability of neural prosthetic control signals from silicon cortical arrays in rhesus macaque motor cortex. Journal of Neural Engineering 8, 045005. URL: https://dx.doi.org/10.1088/1741-2560/8/4/045005. doi: 10.1088/1741-2560/8/4/045005.

9. Prasad, A., and Sanchez, J.C. (2012). Quantifying long-term microelectrode array functionality using chronic in vivo impedance testing. J Neural Eng 9, 026028.

10. Rousche, P.J., and Normann, R.A. (1998). Chronic recording capability of the utah intracortical electrode array in cat sensory cortex. Journal of Neuroscience Methods 82, 1–15. URL: https://www.sciencedirect.com/science/article/pii/S0165027098000314. doi: 10.1016/S0165-0270(98)00031-4.

11. Ludwig, K.A., Uram, J.D., Yang, J., Martin, D.C., and Kipke, D.R. (2006). Chronic neural recordings using silicon microelectrode arrays electrochemically deposited with a poly(3,4-ethylenedioxythiophene) (pedot) film*. Journal of Neural Engineering 3, 59. URL: https://dx.doi.org/10.1088/1741-2560/3/1/007. doi: 10.1088/1741-2560/3/1/007.

12. Robinson, D. (1968). The electrical properties of metal microelectrodes. Proceedings of the IEEE 56, 1065–1071. doi: 10.1109/PROC.1968.6458.

13. Blau, A. (2013). Cell adhesion promotion strategies for signal transduction enhancement in microelectrode array in vitro electrophysiology: An introductory overview and critical discussion. Current Opinion in Colloid Interface Science 18, 481–492. URL: https://www.sciencedirect.com/science/article/pii/S1359029413000976. doi: 10.1016/j.cocis.2013.07.005.

14. Wheeler, B.C., and Brewer, G.J. (2010). Designing neural networks in culture. Proceedings of the IEEE 98, 398–406. doi: 10.1109/JPROC.2009.2039029.

15. Ludwig, K.A., Miriani, R.M., Langhals, N.B., Joseph, M.D., Anderson, D.J., and Kipke, D.R. (2009). Using a common average reference to improve cortical neuron recordings from microelectrode arrays. Journal of Neurophysiology 101, 1679–1689. URL: https://doi.org/10.1152/jn.90989.2008. doi: 10.1152/jn.90989.2008. arXiv:https://doi.org/10.1152/jn.90989.2008. PMID: 19109453.

16. Eduardo, E., and Burke, D. (1988). The optimal recording electrode configuration for compound sensory action potentials. Journal of Neurology, Neurosurgery & Psychiatry 51, 684–687. URL: https://jnnp.bmj.com/content/51/5/684. doi: 10.1136/jnnp.51.5.684. arXiv:https://jnnp.bmj.com/content/51/5/684.full.pdf.

17. Horch, K.W., and Dhillon, G.S. (2004). Neuroprosthetics Theory and Practice. World Scientific Publishing Co. Pte. Ltd.

18. Swinney, K.R., and Wikswo, J.P., Jr (1980). A calculation of the magnetic field of a nerve action potential. Biophys. J. 32, 719–731.

19. Malmivuo, J., and Plonsey, R. (1995). Bioelectromagnetism: principles and applications of bioelectric and biomagnetic fields. New York: Oxford University Press.

20. Caruso, L., Wunderle, T., Lewis, C.M., Valadeiro, J., Trauchessec, V., Trejo Rosillo, J., Amaral, J.P., Ni, J., Jendritza, P., Fermon, C., Cardoso, S., Freitas, P.P., Fries, P., and Pannetier-Lecoeur, M. (2017). In vivo magnetic recording of neuronal activity. Neuron 95, 1283–1291.e4. URL: https://www.sciencedirect.com/science/article/pii/S0896627317307031. doi: 10.1016/j.neuron.2017.08.012.

21. Lei, Z.Q., Li, G.J., Egelhoff, W.F., Lai, P.T., and Pong, P.W.T. (2011). Review of noise sources in magnetic tunnel junction sensors. IEEE Transactions on Magnetics 47, 602–612. doi: 10.1109/TMAG.2010.2100814.

22. Barbieri, F., Trauchessec, V., Caruso, L., Trejo-Rosillo, J., Telenczuk, B., Paul, E., Bal, T., Destexhe, A., Fermon, C., Pannetier-Lecoeur, M., and Ouanounou, G. (2016). Local recording of biological magnetic fields using giant magneto resistance-based microprobes. Scientific Reports 6, 39330. URL: https://doi.org/10.1038/srep39330. doi: 10.1038/srep39330.

23. Barry, J.F., Turner, M.J., Schloss, J.M., Glenn, D.R., Song, Y., Lukin, M.D., Park, H., and Walsworth, R.L. (2016). Optical magnetic detection of single-neuron action potentials using quantum defects in diamond. Proceedings of the National Academy of Sciences 113, 14133–14138. URL: https://www.pnas.org/doi/abs/10.1073/pnas.1601513113. doi: 10.1073/pnas.1601513113. arXiv:https://www.pnas.org/doi/pdf/10.1073/pnas.1601513113.

24. Weinberg, A.M. (1942). Green’s functions in biological potential problems. The bulletin of mathematical biophysics 4, 107–115. doi: 10.1007/bf02477940.

25. Plonsey, R. (1964). Volume Conductor Fields of Action Currents. Biophysical journal 4, 317–328. doi: 10.1016/s0006-3495(64)86785-0.

26. Plonsey, R. (1965). An Extension of the Solid Angle Potential Formulation for an Active Cell. Biophysical journal 5, 663–667. doi: 10.1016/s0006-3495(65)86744-3.

27. Geselowitz, D.B. (1967). On Bioelectric Potentials in an Inhomogeneous Volume Conductor. Biophysical journal 7, 1–11. doi: 10.1016/s0006-3495(67)86571-8.

28. Stratton, J.A. (2007). Electromagnetic theory. IEEE Press series on electromagnetic wave theory reissued ed ed.. Piscataway, NJ: IEEE Press [u.a.]. ISBN 978-0-470-13153-4.

29. Pettersen, K.H., and Einevoll, G.T. (2008). Amplitude variability and extracellular low-pass filtering of neuronal spikes. Biophysical Journal 94, 784–802. URL: https://www.sciencedirect.com/science/article/pii/S0006349508706799. doi: 10.1529/biophysj.107.111179.

30. Buzsáki, G., Anastassiou, C.A., and Koch, C. (2012). The origin of extracellular fields and currents — eeg, ecog, lfp and spikes. Nature Reviews Neuroscience 13, 407–420. URL: https://doi.org/10.1038/nrn3241. doi: 10.1038/nrn3241.

31. Rall, W. (1962). Electrophysiology of a dendritic neuron model. Biophys J 2, 145–167.

32. Moffitt, M.A., and McIntyre, C.C. (2005). Model-based analysis of cortical recording with silicon microelectrodes. Clinical Neurophysiology 116, 2240–2250. URL: https://www.sciencedirect.com/science/article/pii/S1388245705002257. doi: 10.1016/j.clinph.2005.05.018.

33. Jackson, J.D. (2009). Classical electrodynamics. 3. ed., [nachdr.] ed.. Hoboken, NY: Wiley. ISBN 978-0-471-30932-1.

34. Clark, J., and Plonsey, R. (1966). A Mathematical Evaluation of the Core Conductor Model. Biophysical journal 6, 95–112. doi: 10.1016/s0006-3495(66)86642-0.

35. Geselowitz, D.B. (1966). Comment on the Core Conductor Model. Biophysical journal 6, 691–692. doi: 10.1016/s0006-3495(66)86687-0.

36. Clark, J., and Plonsey, R. (1968). The Extracellular Potential Field of the Single Active Nerve Fiber in a Volume Conductor. Biophysical journal 8, 842–864. doi: 10.1016/s0006-3495(68)86524-5.

37. Carnevale, N.T., and Hines, M.L. (2006). The NEURON Book. Cambridge University Press.

38. Hagen, E., Næss, S., Ness, T.V., and Einevoll, G.T. (2018). Multimodal modeling of neural network activity: Computing lfp, ecog, eeg, and meg signals with lfpy 2.0. Frontiers in Neuroinformatics 12. URL: https://www.frontiersin.org/articles/10.3389/fninf.2018.00092. doi: 10.3389/fninf.2018.00092.

39. Markram, H., Muller, E., Ramaswamy, S., Reimann, M., Abdellah, M., Sanchez, C., Ailamaki, A., Alonso-Nanclares, L., Antille, N., Arsever, S., Kahou, G., Berger, T., Bilgili, A., Buncic, N., Chalimourda, A., Chindemi, G., Courcol, J.D., Delalondre, F., Delattre, V., Druckmann, S., Dumusc, R., Dynes, J., Eilemann, S., Gal, E., Gevaert, M., Ghobril, J.P., Gidon, A., Graham, J., Gupta, A., Haenel, V., Hay, E., Heinis, T., Hernando, J., Hines, M., Kanari, L., Keller, D., Kenyon, J., Khazen, G., Kim, Y., King, J., Kisvarday, Z., Kumbhar, P., Lasserre, S., Le Bé, J.V., Magalhães, B., Merchán-Pérez, A., Meystre, J., Morrice, B., Muller, J., Muñoz-Céspedes, A., Muralidhar, S., Muthurasa, K., Nachbaur, D., Newton, T., Nolte, M., Ovcharenko, A., Palacios, J., Pastor, L., Perin, R., Ranjan, R., Riachi, I., Rodríguez, J.R., Riquelme, J., Rössert, C., Sfyrakis, K., Shi, Y., Shillcock, J., Silberberg, G., Silva, R., Tauheed, F., Telefont, M., Toledo-Rodriguez, M., Tränkler, T., Van Geit, W., Díaz, J., Walker, R., Wang, Y., Zaninetta, S., DeFelipe, J., Hill, S., Segev, I., and Schürmann, F. (2015). Reconstruction and simulation of neocortical microcircuitry. Cell 163, 456–492. URL: https://doi.org/10.1016/j.cell.2015.09.029. doi: 10.1016/j.cell.2015.09.029.

40. Goodman, J.W. (2005). Introduction to Fourier Optics. 3rd ed. Roberts and Company Publishers.

41. Muratore, D.G., Tandon, P., Wootters, M., Chichilnisky, E.J., Mitra, S., and Murmann, B. (2019). A data-compressive wired-or readout for massively parallel neural recording. IEEE Transactions on Biomedical Circuits and Systems 13, 1128–1140. doi: 10.1109/TBCAS.2019.2935468.

42. Tse, D., and Viswanath, P. (2005). Fundamentals of Wireless Communication. 1 ed.. Cambridge University Press. ISBN 978-0-521-84527-4 978-0-511-80721-3. URL: https://www.cambridge.org/core/product/identifier/9780511807213/type/book. doi: 10.1017/CBO9780511807213.

43. Verdú, S. (1998). Multiuser detection. Cambridge: Cambridge University Press.

44. Ito, D., Tamate, H., Nagayama, M., Uchida, T., Kudoh, S., and Gohara, K. (2010). Minimum neuron density for synchronized bursts in a rat cortical culture on multi-electrode arrays. Neuroscience 171, 50–61. URL: https://www.sciencedirect.com/science/article/pii/S0306452210011668. doi: 10.1016/j.neuroscience.2010.08.038.

45. Rey, H.G., Pedreira, C., and Quian Quiroga, R. (2015). Past, present and future of spike sorting techniques. Brain Research Bulletin 119, 106–117. URL: https://www.sciencedirect.com/science/article/pii/S0361923015000684. doi: 10.1016/j.brainresbull.2015.04.007. Advances in electrophysiological data analysis.

46. Pachitariu, M., Steinmetz, N.A., Kadir, S.N., Carandini, M., and Harris, K.D. (2016). Fast and accurate spike sorting of high-channel count probes with kilosort. In D. Lee, M. Sugiyama, U. Luxburg, I. Guyon, and R. Garnett, eds. Advances in Neural Information Processing Systems vol. 29. Curran Associates, Inc. URL: https://proceedings.neurips.cc/paper_files/paper/2016/file/1145a30ff80745b56fb0cecf65305017-Paper.pdf.

47. Buccino, A.P., Hurwitz, C.L., Garcia, S., Magland, J., Siegle, J.H., Hurwitz, R., and Hennig, M.H. (2020). Spikeinterface, a unified framework for spike sorting. Elife 9, e61834.

48. Rossant, C., Kadir, S.N., Goodman, D.F.M., Schulman, J., Hunter, M.L.D., Saleem, A.B., Grosmark, A., Belluscio, M., Denfield, G.H., Ecker, A.S., Tolias, A.S., Solomon, S., Buzsáki, G., Carandini, M., and Harris, K.D. (2016). Spike sorting for large, dense electrode arrays. Nature Neuroscience 19, 634–641. URL: https://doi.org/10.1038/nn.4268. doi: 10.1038/nn.4268.

49. Buzsáki, G. (2004). Large-scale recording of neuronal ensembles. Nature Neuroscience 7, 446–451. URL: https://doi.org/10.1038/nn1233. doi: 10.1038/nn1233.

50. Buccino, A.P., Yuan, X., Emmenegger, V., Xue, X., Gänswein, T., and Hierlemann, A. (2022). An automated method for precise axon reconstruction from recordings of high-density micro-electrode arrays. Journal of Neural Engineering 19, 026026. URL: https://dx.doi.org/10.1088/1741-2552/ac59a2. doi: 10.1088/1741-2552/ac59a2.

51. Bullmann, T., Radivojevic, M., Huber, S.T., Deligkaris, K., Hierlemann, A., and Frey, U. (2019). Large-scale mapping of axonal arbors using high-density microelectrode arrays. Frontiers in Cellular Neuroscience 13. URL: https://www.frontiersin.org/journals/cellular-neuroscience/articles/10.3389/fncel.2019.00404. doi: 10.3389/fncel.2019.00404.

52. Bakkum, D.J., Frey, U., Radivojevic, M., Russell, T.L., Müller, J., Fiscella, M., Takahashi, H., and Hierlemann, A. (2013). Tracking axonal action potential propagation on a high-density microelectrode array across hundreds of sites. Nature Communications 4, 2181. URL: https://doi.org/10.1038/ncomms3181. doi: 10.1038/ncomms3181.

53. Cortes, C., and Vapnik, V. (1995). Support-vector networks. Machine Learning 20, 273–297. URL: https://doi.org/10.1007/BF00994018. doi: 10.1007/BF00994018.

54. Lee, Y.C., Chao, C.T., Li, L.C., Suen, Y.W., Horng, L., Wu, T.H., Chang, C.R., and Wu, J.C. (2015). Magnetic tunnel junction based out-of-plane field sensor with perpendicular magnetic anisotropy in reference layer. Journal of Applied Physics 117, 17A320.

55. Holt, G.R., and Koch, C. (1999). Electrical interactions via the extracellular potential near cell bodies. J. Comput. Neurosci. 6, 169–184.

56. Buccino, A.P., and Einevoll, G.T. (2021). Mearec: A fast and customizable testbench simulator for ground-truth extracellular spiking activity. Neuroinformatics 19, 185–204. URL: https://doi.org/10.1007/s12021-020-09467-7. doi: 10.1007/s12021-020-09467-7.

57. Griffiths,., David J. (David Jeffery) ([2013]). Introduction to electrodynamics. Fourth edition. Boston : Pearson, [2013] ©2013. URL: https://search.library.wisc.edu/catalog/9910134691602121 includes bibliographical references and index.

58. Fan, B., Wolfrum, B., and Robinson, J.T. (2021). Impedance scaling for gold and platinum microelectrodes. Journal of Neural Engineering 18, 056025. URL: https://dx.doi.org/10.1088/1741-2552/ac20e5. doi: 10.1088/1741-2552/ac20e5.

59. Wang, A., Jung, D., Lee, D., and Wang, H. (2021). Impedance characterization and modeling of subcellular to micro-sized electrodes with varying materials and pedot:pss coating for bioelectrical interfaces. ACS Applied Electronic Materials 3, 5226–5239. URL: https://doi.org/10.1021/acsaelm.1c00687. doi: 10.1021/acsaelm.1c00687.

60. Ye, Z., Shelton, A.M., Shaker, J.R., Boussard, J., Colonell, J., Manavi, S., Chen, S., Windolf, C., Hurwitz, C., Namima, T., Pedraja, F., Weiss, S., Raducanu, B., Ness, T.V., Einevoll, G.T., Laurent, G., Sawtell, N.B., Bair, W., Pasupathy, A., Lopez, C.M., Dutta, B., Paninski, L., Siegle, J.H., Koch, C., Olsen, S.R., Harris, T.D., and Steinmetz, N.A. (2023). Ultra-high density electrodes improve detection, yield, and cell type specificity of brain recordings. bioRxiv. URL: https://www.biorxiv.org/content/early/2023/08/25/2023.08.23.554527. doi: 10.1101/2023.08.23.554527. arXiv:https://www.biorxiv.org/content/early/2023/08/25/2023.08.23.554527.full.pd.f

61. Parashar, M., Saha, K., and Bandyopadhyay, S. (2020). Axon hillock currents enable single-neuron-resolved 3d reconstruction using diamond nitrogen-vacancy magnetometry. Communications Physics 3, 174. URL: https://doi.org/10.1038/s42005-020-00439-6. doi: 10.1038/s42005-020-00439-6.

