## Supplemental Proofs and Figure 1 for "Spatial Richness of Neural Magnetic Fields"

**Supplemental Material**
Ziad Ali[1,2*] and Ada S. Y. Poon[1,3*]

[1]Stanford University, Stanford, CA, USA
[2]Lead Contact
[3]Senior Author

### Proof of Equation (2) in Main Text

By choosing $\mathbf{P} = \mathbf{H}$ and $\mathbf{Q} = \mathbf{c}/|\mathbf{r} - \mathbf{r}'|$ for any constant vector $\mathbf{c}$ in the vector form of Green's theorem (Equation (1) in main text), we obtain

$$\begin{aligned} &\mathbf{c} \cdot \int_{\mathcal{V}} \frac{\nabla' \times \nabla' \times \mathbf{H}(\mathbf{r}')}{|\mathbf{r} - \mathbf{r}'|} \, d^3r' - \int_{\mathcal{V}} \mathbf{H}(\mathbf{r}') \cdot \nabla' \times \nabla' \times \frac{\mathbf{c}}{|\mathbf{r} - \mathbf{r}'|} \, d^3r' \\ &= \oint_{\mathcal{S}} \mathbf{H}(\mathbf{r}') \times \nabla' \times \frac{\mathbf{c}}{|\mathbf{r} - \mathbf{r}'|} \cdot \hat{\mathbf{n}}' \, da' - \oint_{\mathcal{S}} \left[ \frac{\mathbf{c}}{|\mathbf{r} - \mathbf{r}'|} \times \nabla' \times \mathbf{H}(\mathbf{r}') \right] \cdot \hat{\mathbf{n}}' \, da'. \end{aligned} \tag{1}$$

In the second term, we have

$$\nabla' \times \nabla' \times \frac{\mathbf{c}}{|\mathbf{r} - \mathbf{r}'|} = \nabla' \left( \nabla' \cdot \frac{\mathbf{c}}{|\mathbf{r} - \mathbf{r}'|} \right) - \nabla'^2 \frac{\mathbf{c}}{|\mathbf{r} - \mathbf{r}'|} = \nabla' \left( \nabla' \cdot \frac{\mathbf{c}}{|\mathbf{r} - \mathbf{r}'|} \right) + 4\pi \mathbf{c} \, \delta(\mathbf{r} - \mathbf{r}') \tag{2a}$$

and

$$\mathbf{H}(\mathbf{r}') \cdot \nabla' \left( \nabla' \cdot \frac{\mathbf{c}}{|\mathbf{r} - \mathbf{r}'|} \right) = \nabla' \cdot \left[ \mathbf{H}(\mathbf{r}') \nabla' \cdot \frac{\mathbf{c}}{|\mathbf{r} - \mathbf{r}'|} \right] = \nabla' \cdot \left[ \left( \mathbf{c} \cdot \nabla' \frac{1}{|\mathbf{r} - \mathbf{r}'|} \right) \mathbf{H}(\mathbf{r}') \right]. \tag{3}$$

Substituting into the second term yields

$$\int_{\mathcal{V}} \mathbf{H}(\mathbf{r}') \cdot \nabla' \times \nabla' \times \frac{\mathbf{c}}{|\mathbf{r} - \mathbf{r}'|} \, d^3r' = \int_{\mathcal{S}} \left( \mathbf{c} \cdot \nabla' \frac{1}{|\mathbf{r} - \mathbf{r}'|} \right) \mathbf{H}(\mathbf{r}') \cdot \hat{\mathbf{n}}' \, da' + 4\pi \int_{\mathcal{V}} \mathbf{H}(\mathbf{r}') \cdot \mathbf{c} \, \delta(|\mathbf{r} - \mathbf{r}'|) \, d^3r' \tag{4a}$$

$$= \mathbf{c} \cdot \left\{ \int_{\mathcal{S}} \nabla' \frac{1}{|\mathbf{r} - \mathbf{r}'|} \left[ \mathbf{H}(\mathbf{r}') \cdot \hat{\mathbf{n}}' \right] da' + 4\pi \int_{\mathcal{V}} \mathbf{H}(\mathbf{r}') \, \delta(|\mathbf{r} - \mathbf{r}'|) \, d^3r' \right\}. \tag{4b}$$

In the third term, we have

$$\begin{aligned} \mathbf{H}(\mathbf{r}') \times \nabla' \times \frac{\mathbf{c}}{|\mathbf{r} - \mathbf{r}'|} \cdot \hat{\mathbf{n}}' &= -\nabla' \times \frac{\mathbf{c}}{|\mathbf{r} - \mathbf{r}'|} \cdot \left[ \mathbf{H}(\mathbf{r}') \times \hat{\mathbf{n}}' \right] && \text{(5a)} \\ &= \left( \mathbf{c} \times \nabla' \frac{1}{|\mathbf{r} - \mathbf{r}'|} \right) \cdot \left[ \mathbf{H}(\mathbf{r}') \times \hat{\mathbf{n}}' \right] && \text{(5b)} \\ &= \mathbf{c} \cdot \left\{ \nabla' \frac{1}{|\mathbf{r} - \mathbf{r}'|} \times \left[ \mathbf{H}(\mathbf{r}') \times \hat{\mathbf{n}}' \right] \right\}. && \text{(5c)} \end{aligned}$$

Finally, the integrand in the fourth term can be written as

$$\left[ \frac{\mathbf{c}}{|\mathbf{r} - \mathbf{r}'|} \times \nabla' \times \mathbf{H}(\mathbf{r}') \right] \cdot \hat{\mathbf{n}}' = \mathbf{c} \cdot \left[ \frac{\nabla' \times \mathbf{H}(\mathbf{r}')}{|\mathbf{r} - \mathbf{r}'|} \times \hat{\mathbf{n}}' \right]. \tag{6}$$

Putting together, we obtain Equation (2) in the main text.

### Proof of Equation (11) in Main Text

The longitudinal and the transmembrane components of the intracellular surface current density are given by

$$i^l_{is,z}(z) = -\sigma_i \frac{\partial \phi_i(\mathbf{r})}{\partial z}\bigg|_{\rho=a} \quad \text{and} \quad i^t_{is,\rho}(z) = -\sigma_i \frac{\partial \phi_i(\mathbf{r})}{\partial \rho}\bigg|_{\rho=a}, \tag{7}$$

respectively. Applying Fourier transform along $z$, we obtain

$$I^l_{is,z}(k) = -\sigma_i \cdot ik\Phi_i(a,k) \quad \text{and} \quad I^t_{is,\rho}(k) = -\sigma_i \frac{\partial \Phi_i(\rho,k)}{\partial \rho}\bigg|_{\rho=a}, \tag{8}$$

respectively. From Equation (10) in the main text, we have $\Phi_i(\rho,k) \approx I_0(|k|\rho)\,\Phi_m(k)$. Therefore, we obtain

$$I^l_{is,z}(k) \approx -jk\sigma_i I_0(|k|a)\,\Phi_m(k) \quad \text{and} \quad I^t_{is,\rho}(k) \approx -\sigma_i |k| I_0'(|k|a)\,\Phi_m(k), \tag{9}$$

As $|k|a \ll 1$, $I_0(|k|a) \approx 1$ and $I_0'(|k|a) = I_1(|k|a) \approx \frac{|k|a}{2}$. This yields Equation (11) in the main text.

### Proof of Equation (13) in Main Text

From Equation (5a) in the Main Text, the extracellular magnetic field is given by

$$\mathbf{B}_e(\mathbf{r}) \approx \frac{\mu_0}{4\pi} \oint_{\mathcal{S}} \frac{-\sigma_i \nabla' \phi_i(\mathbf{r}') \times \hat{\mathbf{n}}'}{|\mathbf{r}-\mathbf{r}'|}\, da' \tag{10a}$$

$$= -\frac{\sigma_i \mu_0}{4\pi} \oint_{\mathcal{S}} \frac{1}{|\mathbf{r}-\mathbf{r}'|} \frac{\partial \phi_i(\rho', z')}{\partial z'} \hat{\boldsymbol{\phi}}'\, da'. \tag{10b}$$

We decompose the scalar Green's function, $\frac{1}{|\mathbf{r}-\mathbf{r}'|}$, into a summation of orthogonal functions in the cylindrical coordinates:

$$\frac{1}{|\mathbf{r}-\mathbf{r}'|} = \sum_{m=-\infty}^{\infty} \int_{-\infty}^{\infty} \psi(m,k;\mathbf{r}) \tilde{\psi}(m,k;\mathbf{r}') dk \tag{11}$$

where for $\rho > \rho'$,

$$\psi(m,k;\mathbf{r}) = \frac{1}{\sqrt{\pi}} e^{jm\phi} e^{jkz} K_m(|k|\rho) \tag{12a}$$

$$\tilde{\psi}(m,k;\mathbf{r}) = \frac{1}{\sqrt{\pi}} e^{-jm\phi} e^{-jkz} I_m(|k|\rho), \tag{12b}$$

and $I_m(\cdot)$ and $K_m(\cdot)$ are modified Bessel functions of the first and second kind of order $m$, respectively. Now, we can solve (10b) by separation of variables:

$$\mathbf{B}_e(\mathbf{r}) \approx -\frac{\sigma_i \mu_0}{4\pi} \sum_{m=-\infty}^{\infty} \int_{-\infty}^{\infty} \psi(m,k;\mathbf{r}) \oint_{\mathcal{S}} \tilde{\psi}(m,k;\mathbf{r}) \frac{\partial \phi_i(\rho', z')}{\partial z'} \hat{\boldsymbol{\phi}}'\, da'\, dk \tag{13a}$$

$$= -\frac{\sigma_i \mu_0}{4\pi} \sum_{m=-\infty}^{\infty} \int_{-\infty}^{\infty} \psi(m,k;\mathbf{r}) I_m(|k|a) \cdot \frac{a}{\sqrt{\pi}} \int_0^{2\pi} e^{-jm\phi'} (-\sin\phi' \hat{\mathbf{x}} + \cos\phi' \hat{\mathbf{y}})\, d\phi'$$

$$\cdot \int \frac{\partial \phi_i(a, z')}{\partial z'} e^{-jkz'}\, dz'\, dk. \tag{13b}$$

Recognizing the followings:

$$\int_0^{2\pi} e^{-jm\phi} \sin\phi \, d\phi = j\pi \left(\delta_{m+1} - \delta_{m-1}\right) \tag{14a}$$

$$\int_0^{2\pi} e^{-jm\phi} \cos\phi \, d\phi = \pi \left(\delta_{m+1} + \delta_{m-1}\right) \tag{14b}$$

$$\int \frac{\partial \phi_i(a,z')}{\partial z'} e^{-jkz'} \, dz' = ik\Phi_i(a,k), \tag{14c}$$

we obtain

$$\mathbf{B}_e(\mathbf{r}) \approx -\hat{\boldsymbol{\phi}} \frac{\sigma_i \mu_0 a}{2\pi} \int_{-\infty}^{\infty} jke^{jkz} K_1(|k|\rho) I_1(|k|a) \Phi_i(a,k) \, dk. \tag{15}$$

From Equation (10) in the Main Text and for $|k|a \ll 1$,

$$\mathbf{B}_e(\mathbf{r}) \approx -\hat{\boldsymbol{\phi}} \frac{j\sigma_i \mu_0 a^2}{4\pi} \int_{-\infty}^{\infty} k|k| K_1(|k|\rho) \Phi_m(a,k) e^{jkz} \, dk \tag{16}$$

The inverse Fourier transform of $k|k|K_1(|k|\rho)$ is $\frac{j3z\rho}{2(z^2+\rho^2)^{5/2}}$ and therefore, we obtain the expression in Equation (13a) in the Main Text.

Following the approach outlined in Refs. 1–3, we obtain the extracellular potential in the Fourier domain:

$$\Phi_e(\rho,k) = -\frac{\frac{K_0(|k|\rho)}{K_0(|k|a)}}{1 + \frac{\sigma_e}{\sigma_i} \frac{K_1(|k|a)}{K_0(|k|a)} \frac{I_0(|k|a)}{I_1(|k|a)}} \Phi_m(k) \tag{17}$$

where $a$ is the radius of the cylindrical axon. For $|k|a \ll 1$,

$$\Phi_e(\rho,k) \approx \frac{\sigma_i}{2\sigma_e} (ka)^2 \ln(|k|a) \frac{K_0(|k|\rho)}{K_0(|k|a)} \Phi_m(k). \tag{18}$$

Its inverse Fourier transform yields $\phi_e(\rho,z)$. Then, we obtain the extracellular electric field from the gradient of $\phi_e(\rho,z)$.

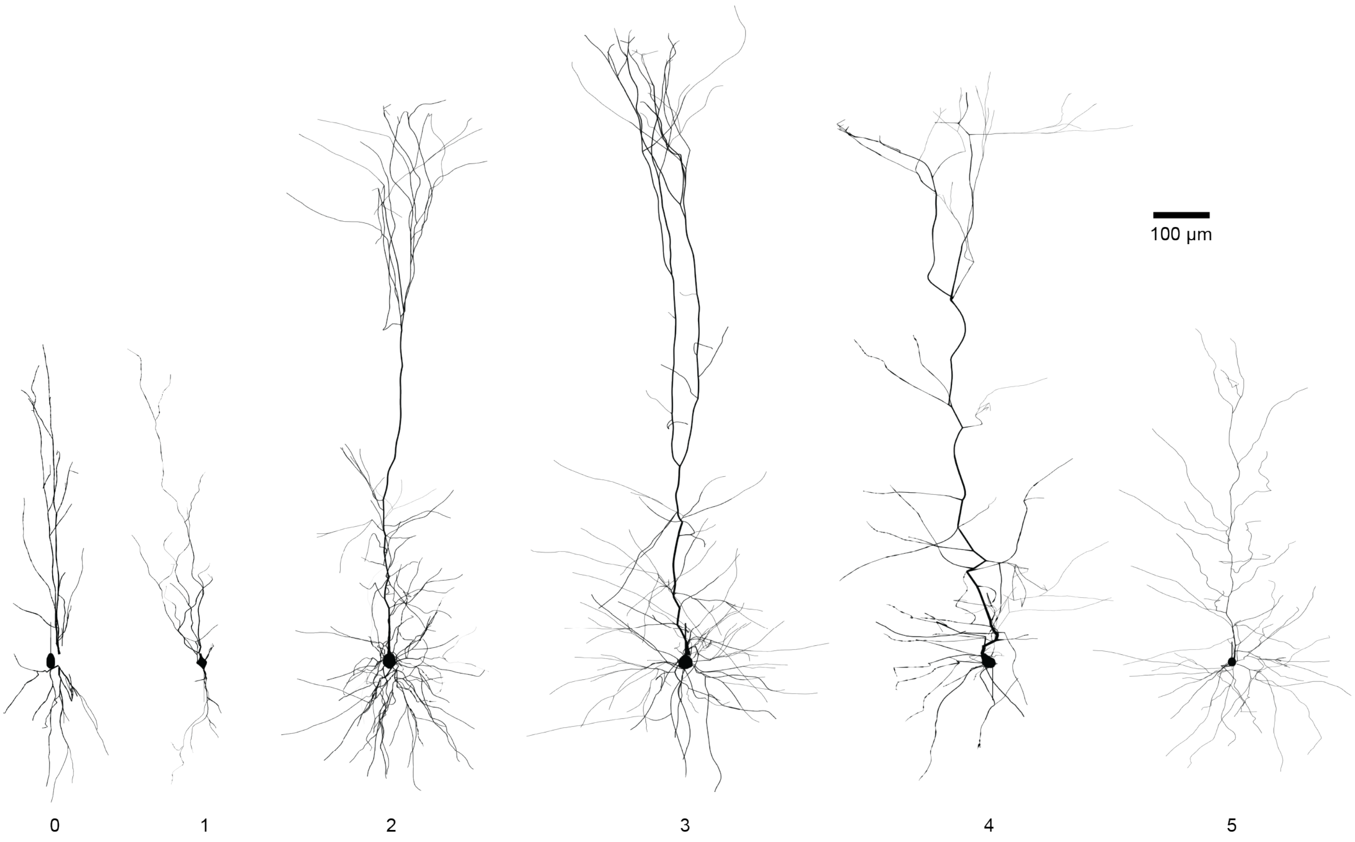


Figure 1: **The six layer V rat somatosensory cortex neuron models ($n = 0 - 5$) used in the study.** Models obtained from Blue Brain Project (BBP)[4]. Model 0: Double bouquet cell (DBC, BBP ID: L5_DBC_bAC217_1). Model 1: Martinotti cell (MC, BBP ID: L5_MC_bAC217_1). Model 2: Thick-tufted pyramidal cell with a late bifurcating apical tuft (TTPC1, BBP ID: L5_TTPC1_cADpyr232_1). Model 3: Thick-tufted pyramidal cell with an early bifurcating apical tuft (TTPC2, BBP ID: L5_TTPC2_cADpyr232_1). Model 4: Slender-tufted pyramidal cell (STPC, BBP ID: L5_STPC_cADpyr232_1). Model 5: Untufted pyramidal cell (UTPC, BBP ID: L5_UTPC_cADpyr232_1).

### References


1. Clark, J., and Plonsey, R. (1966). A Mathematical Evaluation of the Core Conductor Model. Biophysical journal *6*, 95 – 112. doi: 10.1016/s0006-3495(66)86642-0.

2. Geselowitz, D.B. (1966). Comment on the Core Conductor Model. Biophysical journal *6*, 691 – 692. doi: 10.1016/s0006-3495(66)86687-0.

3. Clark, J., and Plonsey, R. (1968). The Extracellular Potential Field of the Single Active Nerve Fiber in a Volume Conductor. Biophysical journal *8*, 842 – 864. doi: 10.1016/s0006-3495(68)86524-5.

4. Ramaswamy, S., Courcol, J.D., Abdellah, M., Adaszewski, S.R., Antille, N., Arsever, S., Atenekeng, G., Bilgili, A., Brukau, Y., Chalimourda, A., Chindemi, G., Delalondre, F., Dumusc, R., Eilemann, S., Gevaert, M.E., Gleeson, P., Graham, J.W., Hernando, J.B., Kanari, L., Katkov, Y., Keller, D., King, J.G., Ranjan, R., Reimann, M.W., Rössert, C., Shi, Y., Shillcock, J.C., Telefont, M., Van Geit, W., Diaz, J.V., Walker, R., Wang, Y., Zaninetta, S.M., DeFelipe, J., Hill, S.L., Muller, J., Segev, I., Schürmann, F., Muller, E.B., and Markram, H. (2015). The neocortical microcircuit collaboration portal: a resource for rat somatosensory cortex. Front. Neural Circuits *9*, 44.
